# Abstraction of reward context facilitates relative reward coding in neural populations of the anterior cingulate cortex

**DOI:** 10.1101/2022.12.10.519901

**Authors:** Jonathan M. Chien, Joni D. Wallis, Erin L. Rich

## Abstract

The anterior cingulate cortex (ACC) is believed to be involved in many cognitive processes, including linking goals to actions and tracking decision-relevant contextual information. ACC neurons robustly encode expected outcomes, but how this relates to putative functions of ACC remains unknown. Here, we approach this question from the perspective of population codes by analyzing neural spiking data in the ventral and dorsal banks of the ACC in monkeys trained to perform a stimulus-motor mapping task. We found that neural populations favor a representational geometry that emphasizes contextual information, while facilitating the independent, abstract representation of multiple task-relevant variables. In addition, trial outcomes were primarily encoded relative to task context, suggesting that the population structures we observe could be a mechanism allowing feedback to be interpreted in a context-dependent manner. Together, our results point to a prominent role for ACC in context-setting and relative interpretation of outcomes, facilitated by abstract, or “untangled,” representations of task variables.

**Author Summary:** The ability to interpret events in light of the current context is a critical facet of higher-order cognition. The anterior cingulate cortex is suggested to be important for tracking information about current contexts, while alternate views hold that its function is more related to the motor system and linking goals to appropriate motor responses. Here, we evaluated these two possibilities by recording anterior cingulate neurons from monkeys performing a stimulus-motor mapping task in which compound cues both defined the current reward context and instructed appropriate motor responses. By analyzing geometric properties of neural population activity, we found that the ACC prioritized context information, representing it as a dominant, abstract concept. Ensuing trial outcomes were then coded relative to these contexts, suggesting an important role for these representations in context-dependent evaluation. Such mechanisms may be critical for the abstract reasoning and generalization characteristic of biological intelligence.

## Introduction

Neurons encoding expected outcomes are commonly found in multiple brain areas, including the orbitofrontal cortex (OFC) and anterior cingulate cortex (ACC) (1 - 7). In OFC, outcome expectations are believed to play a role in choice behavior (8, 9), but in ACC the importance of these signals is less clear. Beyond decision-making, expecting a particular outcome can motivate motor responses and provide context for determining whether a result was better or worse than expected. Different lines of evidence support a role for ACC in each of these processes. On one hand, ACC has been implicated in goal-based motor selection (7, 10), as well as computing decisions from effort or action costs (3, 11, 12). On the other, ACC also tracks contextual information important for accurately predicting and interpreting outcomes, such as richness, temporal proximity, or volatility of rewards in an environment (13 - 16). Here, we aimed to advance our understanding of expectation signals in ACC by determining how neurons code potential reward or punishment in relation to motor responses and outcomes that are eventually received.

An important challenge in interpreting expectation signals in ACC is that single neuron responses are highly heterogeneous, even within the same task paradigm (7, 17, 18). There is increasing evidence that computations arising from such heterogeneity may be best understood by examining population codes (19, 20, 21). Single neurons with heterogeneous and nonlinear responses produce population activity that is high-dimensional, a property that is advantageous for flexibly encoding different combinations of task variables (22, 23). However, when single neuron responses are more linear or not entirely heterogeneous, lower-dimensional structure can be resolved in the population activity. In these cases, investigating representational structure can reveal how information is organized by neural circuits (24, 25). In higher cortical areas, this structure typically represents latent factors that correspond to concepts or variables relevant to the region’s function and are invariant to other features of incoming stimuli. For instance, consistent with a role in object recognition, representations in inferotemporal cortex (IT) distinguish visual images invariant to their rotation or size (26 - 28), and individual IT neurons display tuning to particular visual features, such as curvature, or to particular axes in a “face space,” with insensitivity to other features and axes (29 - 31). Such representations are said to be ‘untangled’ (or ‘disentangled’). Similar conceptual organizations molded by unique functions of brain regions have also been reported in areas of prefrontal cortex (32, 33).

From this perspective, we aimed to determine whether and how latent factors related to expected and received outcomes, as well as motor responses, structure population representations in ACC. To do this, we analyzed neurons recorded from monkeys performing a stimulus-motor mapping task to either earn rewards (positive valence condition) or avoid punishment (negative valence condition). In the task, compound cues signaled both the valence condition and instructed the motor response, allowing us to discern how these variables are represented in neural activity—jointly or as untangled concepts. In addition, trial outcomes could be interpreted in either an absolute (amount of feedback) or relative (better or worse than expected) sense, allowing us to assess how expectations influence outcome coding. We separately analyzed neural responses in the dorsal (dACC) and ventral (vACC) banks of the anterior cingulate sulcus because previous studies have found that these areas differentially encode valence and motor information. Specifically, motor correlates were more common in dACC, and vACC preferentially responded to negative cues and outcomes, whereas dACC included a mix of neurons that respond to positive and negative information (4, 34).

Our results show that both regions of ACC represented the valence of expected outcomes separately from other task information, in an abstract manner that was independent of the particular cues carrying the information. This organization persisted strongly throughout the trial, resulting in non-random structure in the correlation statistics of population activity. When the outcome of the trial was received, ACC primarily encoded the resulting reward or punishment relative to this abstract valence representation, such that the cue’s valence acted as a context for interpreting outcomes. We suggest that this role in setting the context for subsequent outcomes is a key factor encouraging the abstraction of cue valence in this task, and that context-dependent behavior can be facilitated by untangled representations that generalize across particular stimuli.

## Results

To understand how populations of ACC neurons represent expected outcomes, two monkeys were trained to perform an instructed response task by moving a bidirectional joystick to the right or left, depending on an instructional cue (**Figure 1**). Cues were displayed centrally on a computer screen, with two cues instructing rightward responses and two instructing leftward responses. Two cues (one right and one left) also indicated that the monkey would earn a reward for responding correctly and nothing for responding incorrectly, while the other two cues (also one right and one left) indicated that the monkey would receive a punishment for incorrect responses and nothing for correct responses (**Table 1**). Therefore, each cue carried a unique combination of valence (potential reward/punishment) and response (left/right) information. Reward and punishment were delivered by increasing or decreasing the size of a reward bar visible at the bottom of the task screen. The reward bar was cashed in for a proportional amount of fruit juice after every 6 completed trials, so that monkeys were motivated to earn increases and avoid decreases in bar size.

**Table 1.** Feedback types.

| Cue valence | Better outcome | Worse outcome |
| --- | --- | --- |
| Positive | +1 bar increment | 0 bar increment |
| Negative | 0 bar increment | -1 bar increment |

**Figure 1.**
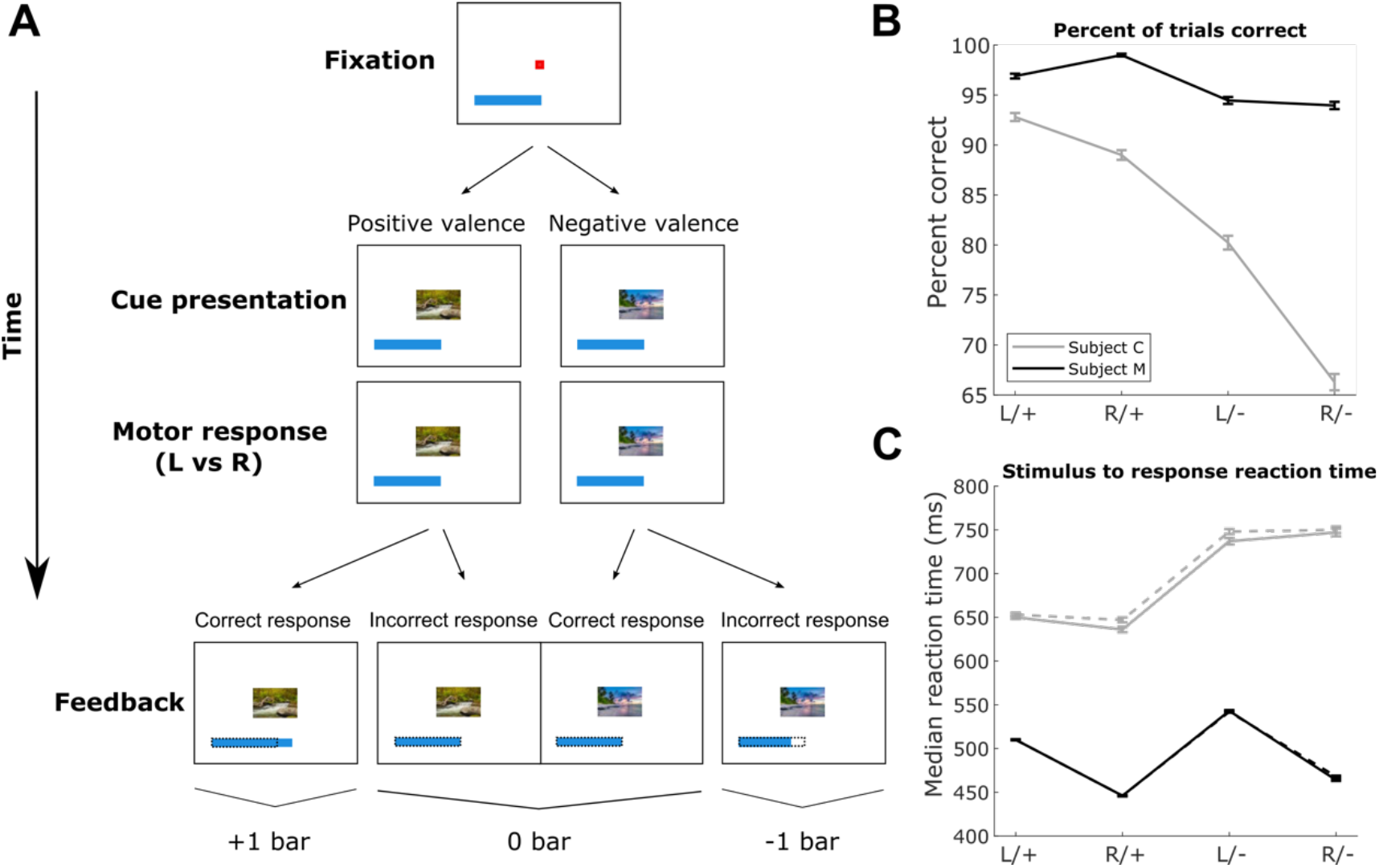
Task and behavioral performance. **(A)** Schematic of the instructed response task. To start a trial, monkeys fixated a central point. Then, one of four familiar pictures appeared in the screen center (Cue presentation). The monkey responded by moving a joystick to the right (R) or left (L) (Motor response). Following a 300 ms delay, the reward bar either did or did not change size, providing feedback for their response. The reward bar was visible throughout the session, and its size carried over to the next trial until 6 trials were completed. At that point, the bar was cashed in for a juice reward proportional to the bar length, and the bar was reset to an initial starting size (not shown). **(B)** Percent of trials of each type completed correctly by each subject. Error bars = +/-SEM across all trials of the same type. (**C**) Median reaction times on trials of each type for each subject. Error bars = +/-standard error of the median across all trials of the same type. Standard errors were computed via bootstrapping. Solid line = correct responses, dashed line = incorrect responses.

Behavior in this task was described in a previous publication (35). Briefly, both subjects performed well, with slightly higher response accuracies and faster response times for positive versus negative cues. Reaction times varied between monkeys, with median values (across all trials from all sessions) of 490 ms and 678 ms for monkeys M and C respectively. Our previous study focused on single neuron responses in subregions of the OFC, and here we assessed similar measures in the dorsal and ventral ACC. We analyzed data from 35 sessions (21 in subject M, 14 in subject C; n = 81 and 64 dACC neurons, and 92 and 70 vACC neurons, in subjects M and C, respectively).

### Valence encoding among single units

To understand how cue information is represented by single neurons in dACC and vACC, we used multiple linear regression (ordinary least squares) to model trial-by-trial firing rates of each neuron (**Methods**). During the cue epoch (101 to 600ms post-cue presentation), we focused on encoding of the cue’s valence (+1/-1) (**Figure 2A & B)** and the response direction it indicated (L/R) (**Figure S2**). Overall, there were more valence selective neurons in dACC than vACC (subject M: χ^2^ = 5.20, p = 0.023; subject C: χ^2^ = 3.20, p = 0.074). Among these, there was a tendency toward different patterns of valence selectivity in each region in subject M, where more dACC neurons were selective for positive than for negative cues, and more vACC neurons were selective for negative than for positive cues, consistent with previous reports (34) (area x selectivity χ^2^ = 3.66, p = 0.056). In subject C, however, this trend was not apparent (χ^2^ = 0.36, p = 0.55). Comparing the average strength of encoding across all neurons and both areas, rather than examining counts of selective neurons, revealed similar results (**Figure S3A & B**). Overall, cue valence was robustly represented in both ACC subregions. While there was some evidence supporting valence selectivity in vACC, these results were inconsistent across subjects.

**Figure 2.**
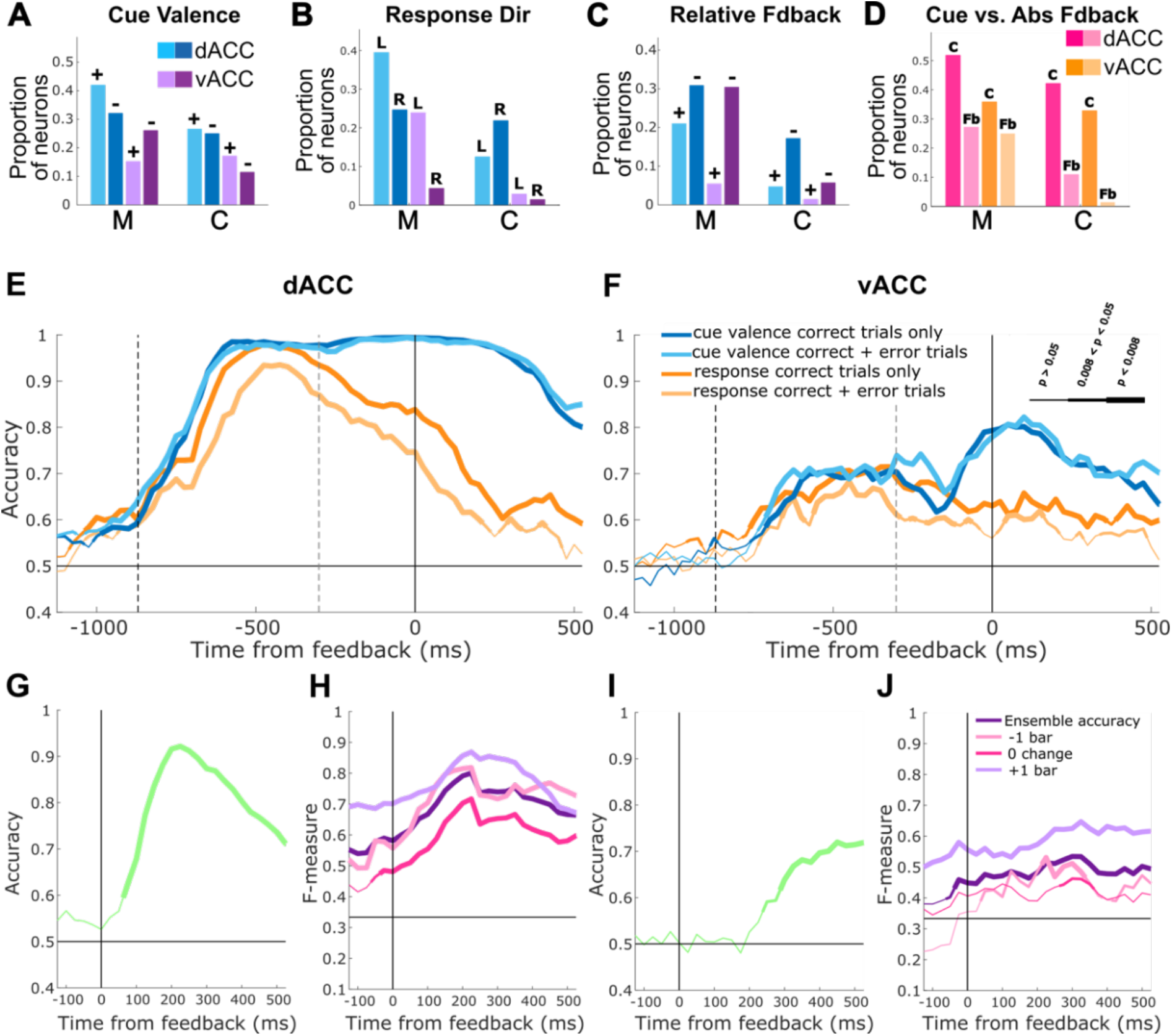
ACC predominantly encodes cue valence and relative feedback. **(A)** Neurons categorized as selective for cue valence during the cue epoch were assigned an encoding preference (positive or negative) based on the mean sign of the regression coefficient during the analysis window. Bars show proportions of valence selective neurons preferring positive (+) and negative (-) cues. **(B)** Same as **A** but showing encoding preferences of neurons selective for response direction during the cue epoch. vACC had more neurons selective for right (R) than left (L) responses in subject M (χ^2^ = 4.33, p = 0.037; post-hoc binomial test on proportion of left-preferring response selective neurons, p = 0.13 and p = 5.34 × 10^−4^ for dACC and vACC respectively) but not subject C (χ^2^ = 1.01, p = 0.31). Note that neurons were recorded from opposite hemispheres in each subject. **(C)** Same as **A** but showing encoding preferences of neurons selective for relative feedback value. **(D)** Each neuron was marked as better explained by cue valence or absolute feedback during the feedback epoch by averaging the difference in *R*^2^ of two respective cue valence and absolute feedback models during the analysis window. Bars show proportions of all neurons in the population whose responses were better explained by cue valence (C) or absolute feedback (Fb) that also demonstrated selectivity for that predictor in the better model. **(E-J)** 10-fold cross-validated decoding accuracy based on population activity in 150 ms bins, stepped by 25 ms and time-locked to feedback presentation (solid vertical line at time = 0 ms). Horizontal lines denote chance performance. **(E)** Decoding accuracy for cue valence and cued response direction in dACC. The black dashed line is the median time of cue presentation across all trials, and the gray dashed line is the median time of the motor response. **(F)** Same as **E** but for vACC. **(G)** Decoding accuracy for relative feedback value in dACC. **(H)** Absolute feedback decoding (ensemble performance and binary one-vs-all performance) in dACC. Note that the (+1)-vs-all and (−1)-vs-all binary classifiers perform above chance prior to feedback, since +1 bar and -1 bar are correlated with positive and negative cue valence. **(I & J)** Same as **G & H**, respectively, but for vACC. Line thickness indicates significance as shown in **F**.

Next, we compared different ways the valence of trial outcomes (feedback) could be coded in this task: as an “absolute” signed magnitude (i.e., +1, 0, -1 for gain, nothing, loss), or as a “relative” value denoting the receipt of the better or worse possible outcome given the preceding cue (**Table 1**). We first assessed relative feedback encoding, which is uncorrelated with the preceding cue, using linear regression. In subject M, there were more vACC neurons selective for worse (negative), compared to better (positive) outcomes (area x valence selectivity χ^2^ =5.72, p =0.017, post-hoc binomial test, p = 6.62 × 10^−5^ in vACC), and no differences in dACC (binomial test p = 0.28) (**Figures 2C & S3**). Although the trend was similar in subject C, there were no statistical differences in valence selectivity by area (χ^2^ = 0.0045, p = 0.95).

We also considered whether neurons encoded the absolute outcome of the trial (gain, nothing, loss). Since these outcomes were correlated with the valence of the preceding cue (**Table 1**), we first fit each neuron with a multiple regression model that included an absolute feedback predictor and then fit the same responses with the same model but with the absolute feedback predictor replaced by a cue valence predictor. We considered a neuron to be better modeled as encoding absolute feedback value if it had a significant regression coefficient for the feedback regressor and a higher R^2^ for the feedback model compared to the cue valence model. Conversely, we considered a neuron to be better described as encoding cue valence at the time of feedback if it had a significant regression coefficient for cue valence, and the cue valence model had a higher R^2^ than the feedback model. We found that absolute feedback encoding was very low before feedback onset, as expected, whereas cue encoding was high (**Figure S3C**). After the onset of feedback, cue valence encoding decreased somewhat but continued to be encoded by ∼20-30% of neurons in each area throughout the remainder of the trial. Overall, in a 500ms epoch following feedback onset, cue valence encoding dominated in both subjects and both areas (binomial test on model preference, subject M dACC p = 0.014, vACC p = 0.12, subject C dACC p = 7.73 × 10^−5^, vACC = 0.0011; area × model preference for subject M χ^2^ = 0.55, p = 0.46, for subject C χ^2^ = 0.42, p = 0.52) (**Figure 2D**). Taken together, single unit analyses showed that the valence of a reinforcement-predicting cue is robustly represented from the time of cue presentation until and even past the time that predicted feedback is obtained.

### Population decoding

Next, we assessed how cue information is coded at the level of neuron populations. To do this, we first decoded the valence and instructed response direction from pseudopopulation activity in each area. Population activity was aligned to feedback presentation and spike counts averaged within 150 ms bins, stepped forward by 25 ms, including 1200 ms preceding feedback to capture presentation of the instructive cue, and 600 ms after feedback. For each time bin, we trained and tested linear support vector machines (SVMs) using 10-fold cross-validation to assess classification of cue valence or response direction (**Methods**). We report accuracies below, but evaluating model performance using the area under the ROC curve (AUROC) yielded qualitatively similar results.

Both cue valence and response direction could be decoded from activity in both regions, with higher accuracies in dACC (**Figure 2E & F**). Interestingly, despite the fact that each cue conveyed both valence and response direction information, the time course of decodability differed for these two variables. For response direction, decoding peaked just before the median time of response and subsequently decayed, whereas valence decoding increased when the cue was presented and persisted past the motor response, through the brief (300 ms) delay preceding feedback, and well beyond the delivery of feedback itself. This effect was present in both regions but was especially evident in dACC. In vACC, decoding of cue valence was less accurate but increased as the time of feedback approached. In addition, the inclusion of error trials, defined as those in which the monkey moved the joystick in the wrong direction, decreased decoding accuracy for response direction but not cue valence (**Figure 2E & F**). This further suggests that the valence and response conveyed by a compound cue are treated as separable types of information, which may have different salience to the animal or play different roles in future behavior.

While cue valence appeared to be persistently encoded into the feedback epoch, another possibility is that the ostensible decodability of cue valence was driven by representations of absolute feedback, which correlates with the valence of the cue. To assess this, we attempted to decode the signed magnitude of feedback received (+1, 0, or -1) from population activity. We used an ensemble classifier that handles multiclass classification by combining the results of constituent binary one-vs-all linear SVMs, where each binary classifier decoded one of the feedback conditions from the others (using 3 one-vs-one constituent classifiers yielded qualitatively similar results). Importantly, for this analysis we balanced the number of trials in which receipt of 0 feedback was preceded by positive and negative cues (**Methods**). Even when these trials are balanced, overall decoding of absolute feedback (combined across the 3 constituent classifiers) should be significantly above chance if neural populations encoded cue valence but *not* absolute feedback, because +1 feedback and -1 feedback always follow positive and negative cues respectively. However, the 0-feedback-vs-all binary classifier should perform poorly, since instances of 0 feedback are always preceded by an equal mix of positive and negative cues. We found that, as expected, overall feedback decoding was above chance in both regions preceding feedback onset, consistent with a correlative contribution of the preceding cue valence to feedback decoding. Critically, the one-vs-all classifier for 0 feedback exhibited above chance performance in dACC but showed minimal significance in vACC (**Figure 2H & J**), indicating that dACC does carry information about absolute feedback, while vACC does not. Rather than the signed magnitude of the feedback itself, vACC encodes the valence of a predictive cue at the time of feedback.

In dACC, this analysis, alongside cue valence decoding, indicated that both cue valence and the signed magnitude of the outcome are encoded at feedback time. If dACC populations only represented the absolute value of feedback, the accuracy of a classifier attempting to decode cue valence should be closer to 0.75, since its performance would be at chance on the 0 feedback trials (which constitute one half of all trials; **Table 2**). However, we observe near perfect decoding accuracy of cue valence throughout the entire trial, beginning shortly after cue onset and lasting well past feedback delivery (**Figure 2E**).

**Table 2.** Task conditions.

| Cue Epoch |  |  |
| --- | --- | --- |
|  | Condition | Abbreviation |
| 1) | Positive cue / Left response | [+cue / L] |
| 2) | Positive cue / Right response | [+cue / R] |
| 3) | Negative cue / Left response | [-cue / L] |
| 4) | Negative cue / Right response | [-cue / R] |
| Feedback Epoch |  |  |
|  | Condition | Abbreviation |
| 1) | Positive cue / Left response / +1 bar increment | [+cue / L / +bar] |
| 2) | Positive cue / Left response / 0 bar increment | [+cue / L / 0bar] |
| 3) | Positive cue / Right response / +1 bar increment | [+cue / R / +bar] |
| 4) | Positive cue / Right response / 0 bar increment | [+cue / R / 0bar] |
| 5) | Negative cue / Left response / 0 bar increment | [-cue / L / 0bar] |
| 6) | Negative cue / Left response / -1 bar increment | [-cue / L / -bar] |
| 7) | Negative cue / Right response / 0 bar increment | [-cue / R / 0bar] |
| 8) | Negative cue / Right response / -1 bar increment | [-cue / R / -bar] |

Taken together, both single unit encoding and population decoding indicate that cue valence is a dominant and persistent signal in both ACC regions. In addition, only dACC reliably carried information about the absolute amount of feedback, whereas both regions processed feedback in a relative sense. One potential reason for cue valence to be held online into the feedback epoch is to serve as a context signal necessary for interpreting outcomes in a relative manner. That is, the +1 and -1 bar outcomes have a clear interpretation, but 0 bar is ambiguous, being the better possible outcome following a negative cue (avoidance of loss) or the worse possible outcome following a positive cue (no gain despite the opportunity). This view would also explain why decoding of cue valence did not decrease compared to decoding of response direction on trials in which the subject erred: 0 bar feedback is interpretable even if response direction is incorrectly encoded, but not if the context signaled by the cue valence is confused (**Figure 2E & F**). In the following analyses, we took more direct approaches to determine the degree to which cue valence signals in ACC may serve a context-relevant function.

### ACC represents valence abstractly

If cue valence sets a context for the trial, it may be useful to represent valence in a consistent format across different cues (e.g., left and right instructing cues), such that positive and negative valences are always distinguished in the population code. In other words, valence information may be abstracted to achieve context-dependent outcome coding, such as relative feedback value. Based on this idea, we assessed the abstract nature of valence encoding by investigating the representational geometry of variables in this task (25). For example, consider the four cues **(Table 2)** represented as points in a vector space where each dimension corresponds to the firing activity of one neuron in the population. If a hyperplane separating representations of [+/R] from [-/R] can also separate [+/L] from [-/L], this “cross-condition” decodability suggests that the concept of positive versus negative valence generalizes across response directions, providing evidence for an abstract representation.

Bernardi *et al*. (25) assessed this operational definition of abstraction by investigating cross-condition decodability within different “dichotomies,” which refer to unique divisions of task conditions into two sets of equal cardinality. For four task conditions during the cue epoch, there are three such dichotomies ([+/R,+/L] vs. [-/R,-/L]; [+/R,-/R] vs. [+/L,-/L]; [+/R,-/L] vs. [-/R,+/L]), with the first two corresponding to identifiable task concepts (cue valence and response direction, respectively). For each of these dichotomies, there are four cross-condition training and testing possibilities. For example, for dichotomy 1 (cue valence), one can train on population responses for [+/R] vs. [-/R] and test whether the solution generalizes to [+/L] vs. [-/L]; train on [+/L] vs. [-/L] and test on [+/R] vs. [-/R]; train on [+/R] vs. [-/L] and test on [+/L] vs. [-/R]; or train on [+/L] vs. [-/R] and test on [+/R] vs. [-/L]. During the feedback epoch, there are eight unique conditions, corresponding to four cues and two potential outcomes each, and thus 35 dichotomies (4 of which are interpretable concepts of cue valence, response direction, relative feedback, and reward bar change versus no change; **Table S1**), with 68 training and testing possibilities for each (**Methods**).

For each dichotomy, the average decoding performance over all training and testing possibilities is termed the “Cross-condition generalization performance” (CCGP) (25). High CCGP indicates robust decodability of a concept regardless of the conditions in the training and testing sets. That is, the concept generalizes over different particular instances that vary along other irrelevant features. Using this approach, we found evidence for abstract representation of cue valence that persisted throughout the entire trial, consistent with our finding that cue valence is persistently represented in ACC. dACC exhibited stronger abstraction of valence information (higher CCGP) than vACC, though both were significant (**Figure 3A & E**). In contrast, the concept of response direction had CCGP near chance levels, consistent with the interpretation that, unlike valence, response direction can but does not need to be abstracted across multiple conditions. Finally, the last dichotomy of cue conditions, which did not have an intuitive interpretation, exhibited below chance CCGP, suggesting that cross-condition generalization in this dimension is compromised in order to achieve abstraction in the valence (and to a lesser extent, response direction) dimension (**Figure S5**).

**Figure 3.**
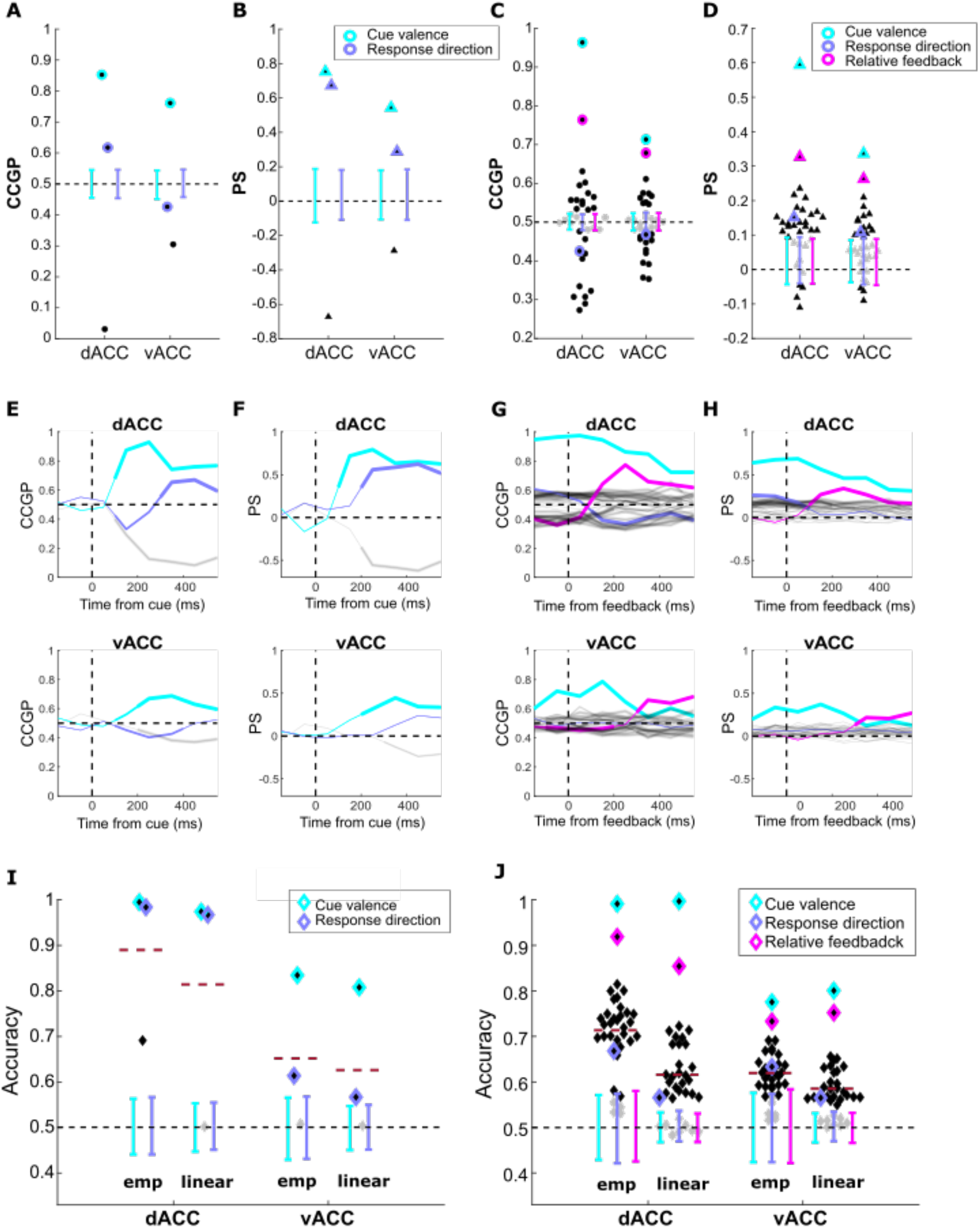
Abstract geometry in ACC. CCGP (**A & C**) and PS (**B & D**) for the stimulus epoch (**A & B**; 3 dichotomies defined over the 4 task conditions) and feedback epoch (**C & D**; 35 dichotomies defined over the 8 task conditions; see **Table S1)**. Each marker in **A-D** represents a dichotomy. Dots denote CCGP, and triangles PS. Black = significance at *α* = 0.05, gray = non-significance. Error bars = 95% CIs for the null distributions of the dichotomy with the matching color. **(E-H)** CCGP (**E & G**) and PS (**F & H**) for dACC (top) and vACC (bottom) during the stimulus epoch (**E & F**; 3 dichotomies) and feedback epoch (**G & H**; 35 dichotomies). Thin lines = non-significance, medium = significance at *α* = 0.05, thick lines = significance at α = 0.01. Gray lines = all remaining dichotomies. **(I)** Decoding performance for each of the 3 cue epoch dichotomies. Red dotted lines show mean performance across all dichotomies (shattering dimensionality). For each brain region, data on the left are decoding results for empirical data, while data on the right are decoding results for the tuned linear (factorized) model. **(J)** Same as **I** but for the feedback epoch.

We also computed the Parallelism Score (PS), an alternative method for assessing population geometry without committing to a particular choice of classifier (25). Briefly, stronger abstraction of a dichotomy should be characterized by more substantial overlap of conditions on the same side of a dichotomy relative to variance across conditions on opposite sides. As such, vector differences linking centroids on opposite sides of the dichotomy should be closer to parallel (**Figure 4F**). To assess this, we calculated cosine similarities of these vector differences in the neural activity space to define the PS, with higher scores indicating stronger abstraction (**Methods**). The PS analysis largely confirmed our CCGP results. Cue valence had the highest scores in both areas, indicating the strongest abstraction (**Figure 3B &F**). Compared to CCGP, evidence for abstraction of response direction was slightly stronger with the PS, but both measures found the remaining undefined dichotomy to be below chance levels, which is consistent with the abstraction of the other two variables. Overall, both salient task variables (cue valence and response direction) are abstracted in both regions during the cue epoch, though more strongly in dACC than vACC, and cue valence was represented more abstractly than response direction.

**Figure 4.**
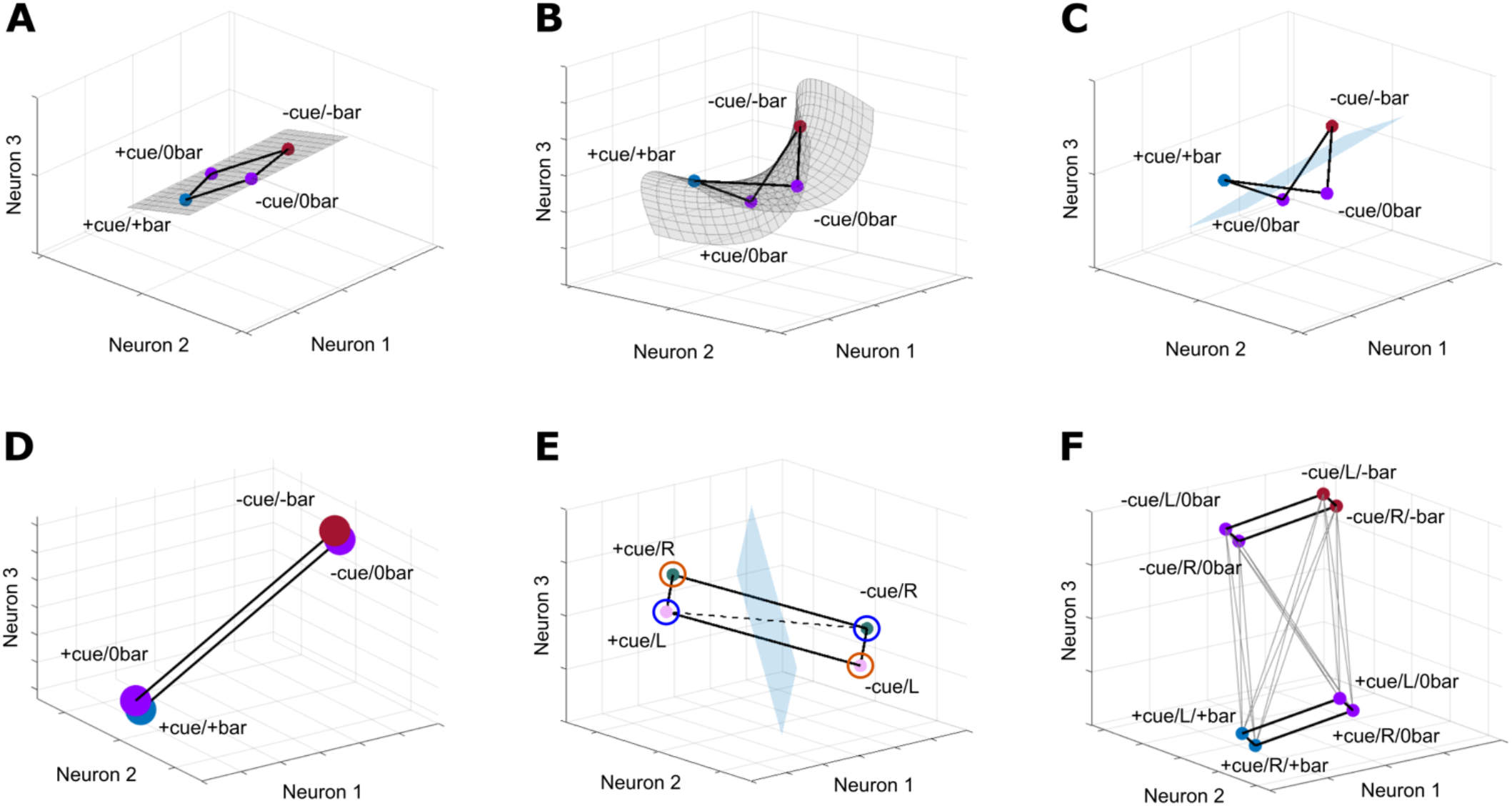
Potential geometries for expected and actual outcome coding. (**A**) Schematic showing neural responses generated as linear combinations of cue valence and relative feedback value. Representations lie on a 2D linear manifold and are captured by a 2D vector subspace of the full 3D neural activity space. **(B**) Similar to **A**, except neural responses incorporate a nonlinear combination of cue valence and relative feedback. Representations thus lie on a 2D nonlinear manifold embedded in a 3D vector subspace (here the full neural activity space). (**C**) The geometry in **B** allows linear separation (blue hyperplane) of the 0 bar vs. bar change conditions necessary for absolute feedback encoding. (**D**) Abstract encoding of cue valence in which instances of each valence condition overlap. Note that encoding cue valence alone, without tracking relative feedback value, could be sufficient to interpret the 0 bar conditions (purple centroids are separated). (**E**) Strong abstraction of cue valence relative to response direction leads to below chance cross-condition decoding for response direction. A decoding hyperplane distinguishing L from R learned from the blue circled training set would actively misclassify direction on the orange circled test set. **(F)** Schematic of the representational geometry in which cue valence, relative feedback, and response direction are abstracted to different degrees. Vector differences (in gray) linking opposite sides of the cue valence dichotomy (bold black squares) are close to parallel, resulting in a higher PS. Vector differences for the relative feedback and response dichotomies would be less and least parallel, respectively, consistent with the level of abstraction observed for each. Note that absolute feedback is not decodable here, as bar change vs. no change conditions are not linearly separable. To achieve this separation, the geometry would have to incorporate a nonlinearity and bend into a fourth embedding dimension, analogously to **B**. Simply moving centroids within the depicted 3D vector space only moves points around on a 3D linear manifold, analogous to that in **A**, and would result in loss of linear separability of another dichotomy. In general, sufficiently weak and specific nonlinearities largely preserve abstract (linear) structure while allowing additional linear separation of select dichotomies.

Next, we used the same methods to determine whether task concepts are abstractly represented during the feedback epoch. Our decoding results showed that cue valence was strongly represented at this time, and here we found that this signal is also abstracted. The highest measures of both CCGP and PS in both regions were for dichotomies corresponding to cue valence (**Figure 3C, D, G, H**). This effect was particularly strong in dACC, where cross-condition decoding accuracy within the feedback epoch approached 1 and dominated all other dichotomies, suggesting that population codes in dACC (and to a lesser extent vACC) strongly prioritize cue valence abstraction. During this epoch, relative feedback was also represented in an abstract fashion in both areas. However, response direction was much different and even displayed below chance CCGP. This indicates that generalizability of response information was de-emphasized by decreasing variance between conditions on opposing sides of a hyperplane separating left versus right. It is likely that this occurred to support the extreme abstraction of cue valence, resulting in classifiers trained under one subset of conditions that actively misclassified conditions from the held-out test set. For example, a decoder tasked with separating the [+/L] and [-/R] cues by response direction will learn a hyperplane more orthogonal to the valence dichotomy and will thus perform below chance on the [-/L] and [+/R] test set, since response direction is preserved but valence is reversed (**Figures 4E, S5 & 8**).

Finally, following Bernardi *et al*., we computed shattering dimensionalities (SD), defined as the mean cross-validated decoding accuracy across all possible dichotomies (25). The goal of this analysis is to better understand the geometries of task representations in high-dimensional neural activity space. For example, consider four conditions defined by all possible combinations of cue valence and relative outcome (**Table 1**). If neurons represent these four conditions as linear combinations of the two variables, then population representations of these conditions would, save for noise, lie on a linear 2D manifold, allowing linear separation of the two dichotomies corresponding to valence and relative outcome but not the last dichotomy corresponding to receipt of feedback (bar change versus no change) (**Figure 4A**). In order to linearly separate the final dichotomy (resulting in a higher SD), the neural representation would have to become nonlinear (lying near a nonlinear 2D manifold), allowing it to bend into a third embedding dimension (**Figure 4B & C;** see **Figure 4F** for generalization to 3 task variables). In other words, a higher SD arises from a more nonlinear population geometry with higher dimensionality as measured by linear methods such as PCA. Experimental and theoretical work has argued that prefrontal cortical neurons tend to have firing rates that encode nonlinear combinations of relevant task variables for this purpose (22, 23). However, abstract neural representations could sacrifice maximal dimensionality by restricting possible geometries to those supporting cross-condition generalization, especially once the task is well-learned (25). We therefore tested the degree to which our recorded neural data exhibits nonlinearities compared to a purely linear model. The SD can be thought of as an overall statistic measuring such nonlinearity, and hence linear separability of conditions in the population geometry, without commenting on the interpretability of any of the dichotomies (**Methods**).

During both the cue and feedback epochs, we found empirical SDs slightly but not substantially higher than that of a model in which neural responses are linear combinations of key variables (cue valence and response direction during the cue epoch, plus relative outcome during the feedback epoch; **Methods**) (**Figures 3I & J)** whose CCGPs match empirical values (**Figure S6**) (cue epoch empirical vs. model SD: 0.89 vs. 0.81 for dACC and 0.65 vs. 0.63 for vACC. Feedback epoch empirical vs. model SD: 0.71 vs. 0.62 for dACC and 0.62 vs. 0.59 for vACC). Larger differences in dACC are consistent with its larger proportion of nonlinear mixed selective neurons compared to vACC (**Figure S4**) and with the decodability of absolute feedback in dACC but not vACC (**Figures 2, 4**). However, compared to SDs in other tasks (25), our data suggest that neural responses in the present task are closer to linear combinations of task variables, with only enough nonlinearity to permit separation of a limited number of other dichotomies (such as bar change vs no change, necessary for absolute feedback encoding).

### Representational similarity and clustering

Since abstraction is driven by clustering of conditions in neural activity space, we reasoned that it should also be reflected in similarity measures, such as correlations and clustering, among conditions. For instance, if cue valence is abstractly represented, two conditions should be highly correlated if they share the same cue valence and anticorrelated if they do not. This can be captured by an ideal correlation pattern (which we term a template) in which all conditions featuring the same cue valence are perfectly correlated, and those with opposite cue valences are perfectly anticorrelated. We performed representational similarity analysis (RSA) (36) to assesses the contributions of six such templates to condition-wise correlation patterns observed in recorded neural data, using linear regression (**Figure 5C**). Empirical correlations were calculated from all pairings of 8 *p*-dimensional vectors of trial-averaged firing rates, where each vector corresponds to mean activity of *p* neurons in one of 8 different task conditions. This was done in sliding 150 ms time bins, stepped at 25 ms. We then calculated the coefficient of partial determination (CPD) for each template regressor to determine its marginal contribution to the empirical condition correlations at each point in time throughout a trial. P-values were computed by comparing results to a null distribution generated by shuffling the empirical correlations (**Methods**).

**Figure 5.**
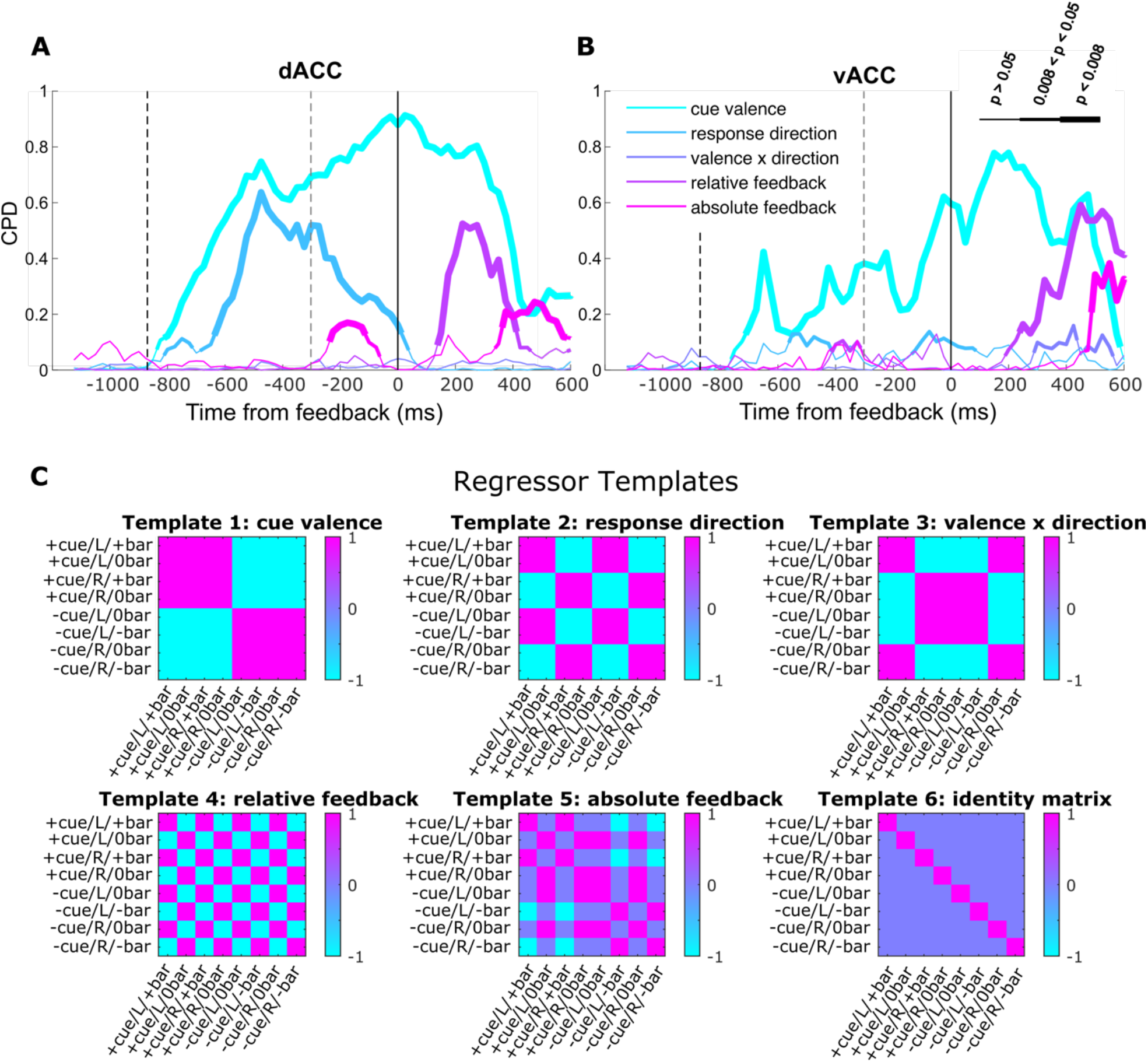
Representational similarity analysis (RSA). Coefficients of partial determination (CPDs) were computed for each template correlation pattern for dACC (**A**) and vACC (**B**). Trials were aligned to the onset of feedback (solid vertical line at time = 0 ms), and black and gray dashed lines indicate median times of cue onset and motor response, respectively. Line thickness denotes level of significance as in the legend based on random permutations of the empirical correlations. (**C**) Templates used in the RSA.

We found that the template matrix for cue valence made the greatest marginal contribution to the model in both regions, consistent with our preceding analyses (**Figure 5**). Representational similarity for cue valence was also stronger in dACC compared to vACC and, in line with decoding results, increased in vACC later in the trial, approaching the time of feedback. After the occurrence of feedback, the most robust and consistent signals in both regions were cue valence and relative feedback, also consistent with our previous analyses, and latencies were again shorter in dACC. Representational similarity of absolute feedback was intermittently present in dACC, and less so in vACC. Together, these results suggest an important role for representations of cue valence that persist during feedback processing.

In addition to cue valence, there was significant representational similarity among conditions featuring the same response direction in dACC (**Figure 5A**). This was predominantly around the time the response was made and decayed by the time of feedback onset. In contrast, vACC showed little representational similarity for response direction (**Figure 5B**), consistent with a previous study that found more motor correlates in dACC (4). This is also consistent with the relative lack of response abstraction in vACC, although this information was decodable above chance levels in the same population. This confirms that significant RSA results are more related to abstraction than to traditional decoding, in that both representational similarity and abstraction require relative overlap between neural responses to stimuli featuring the same value of a task variable (even as other task variables change). Decoding, however, requires no such overlap, but only that population responses be linearly separable. Thus, representational similarity and abstraction imply decodability, but the converse does not hold. Finally, the interaction between valence and response direction accounted for little of the observed correlation patterns throughout the trial in either region.

To more directly study neural population variance across conditions, we used principal components analysis (PCA) to identify latent factors underlying population responses. The activity of each of *p* neurons was averaged both within an analysis window from 101 ms to 600 ms following feedback and across all trials within each of 8 conditions (4 cues x 2 possible outcomes each). Each neuron’s activity was then z-scored across conditions. PCA of the resulting 8 × *p* matrix identified PCs that can be interpreted as canonical responses across conditions. In both areas, the first canonical response was consistent with encoding of cue valence, despite the fact that the data were from the feedback epoch (**Figure 6A-C**). Relative feedback was captured in the second canonical response, confirming the dominance of these two representations in the population. Finally, response direction was extracted with somewhat less fidelity in the third PC, and a nonlinear combination (product) of cue valence and relative feedback, equivalent to the bar change vs no change dichotomy was captured most weakly by the fourth PC.

**Figure 6.**
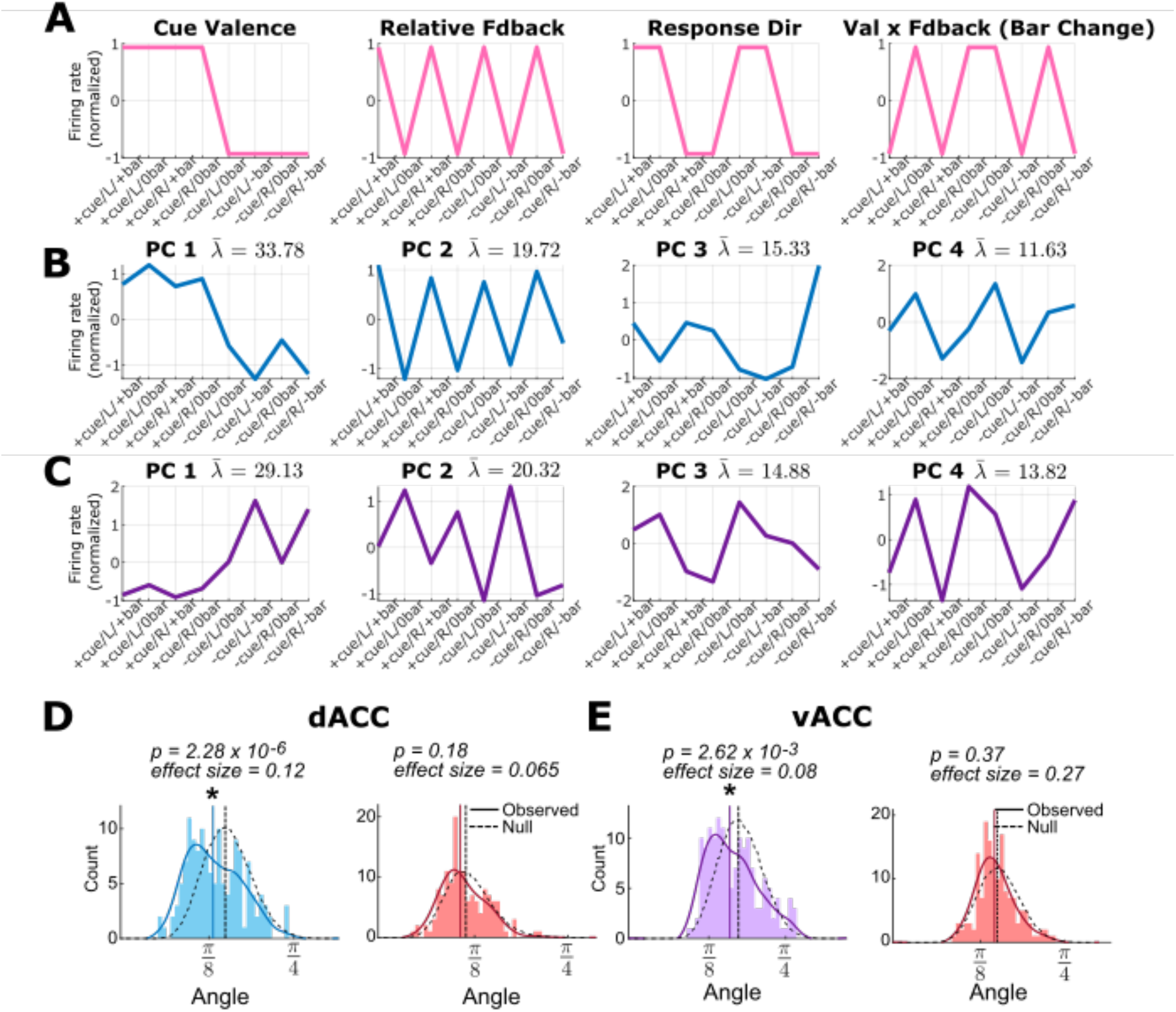
Task variables are nonrandomly and approximately linearly mixed in dACC and vACC. **(A)** Schematic coordinate plots showing canonical response patterns encoding task variables. **(B)** Top 4 standardized PCs from dACC population activity. 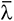 (renormalized eigenvalue) = percent of total variance explained by each PC. **(C)** Same as **B** but for vACC. **(D-E)** Histograms from the ePAIRS analysis showing distributions of mean angles from each neuron’s PCA loading vector to its 3 nearest neighbors (solid lines), compared to a randomly generated null distribution (dashed). Both dACC (**D blue**) and vACC (**E purple**) demonstrated significant clustering. Removing variance associated with only one of the first two PCs reduced but failed to destroy cluster structure in dACC (ePAIRS p = 1.12 × 10^−3^, effect size = 0.09 and p = 2.81 × 10^−4^, effect size = 0.08 upon removal of PC 1 and 2 variance, respectively) but removing variance associated with both did (**D red**). In vACC, removal of variance associated with the first PC was sufficient to destroy cluster structure (**E red**).

PCA also yields loadings that represent the contribution of each standardized PC to the observed standardized activity of each neuron (within the fixed time window). Since PCs can be interpreted as canonical response patterns across different task conditions, often mapping onto a single task variable, we can assess the loadings to understand whether single neurons mix these patterns randomly or in characteristic ways that result in clustering among the loading vectors (37). Recent work in the OFC of macaques and rodents has suggested that prefrontal populations mix task variables nonrandomly, producing clustered responses (38, 39), and theoretical models have been proposed to explain these results (40). To quantify this in ACC, we used the elliptical Projection Angle Index of Response Similarity (ePAIRS) algorithm (37), which assesses the cluster tendency of a set of vectors. The loading weights on the top PCs required to explain 90% of population variance across conditions demonstrated significant cluster tendency in both dACC (**Figure 6D**) and vACC (**Figure 6E**), similar to reports in OFC (38, 39). To determine whether loadings associated with particular PCs drove this clustering, we iteratively removed variance in the population captured by the eigenvector associated with each PC (see **Methods**). Removing the first dimension was sufficient to destroy cluster structure in vACC but not dACC (**Figure 6E**), suggesting that the first PC, which primarily varied with cue valence, drives clustering in vACC. In dACC, removal of the 2 leading PCs was necessary (**Figure 6D**). This suggests that clustering in dACC was also driven mainly by the tendency of neurons to represent cue valence, as well as relative feedback (**Figure 6B & C, S7**).

### Cue valence drives across-condition variance and population dynamics

Finally, in order to summarize our findings and visualize the temporal evolution of task selective information in each region, we projected population activity onto 3D subspaces capturing maximal population variance across time and conditions, as identified by PCA. We used firing rate matrices of size 8*m x p*, with columns corresponding to neurons and rows corresponding to *m* 150 ms time bins (centered from -1000 ms to +600 ms relative to feedback onset, stepped by 25 ms), concatenated across 8 task conditions. The resulting PCs can be interpreted as latent canonical responses across time in each task condition. In dACC, the first and second PCs primarily captured temporal dynamics in the population response, with some selectivity for cue valence at the time of cue presentation (**Figure 7A & B**). The third PC, however, demonstrated clear selectivity for cue valence across the entire trial, most strongly during the feedback epoch. In vACC, this separation of conditions by cue valence was evident in the first PC following the monkeys’ response and leading up to feedback, whereas the third PC revealed the emergence of a separation by cue valence in an orthogonal dimension later in the trial, corresponding to a different pattern of activation over neurons (neural mode). Prior to feedback, dACC also showed a smaller separation according to response direction that was oriented along a different axis than the separation by cue valence. This was most clearly seen in the 3D state space plot (**Figure 7A**), and less so in any individual PC (**Figure 7B**), suggesting that response direction is not entirely captured by any one PC. Similar separation was present but not as dramatic in vACC, consistent with the weaker decoding and abstraction of response direction. Trajectories in both areas remained separated according to cue valence throughout the feedback epoch, at which time they further diverged according to the outcome of the trial to reflect relative feedback. These overall patterns are consistent with a relative reward encoding mechanism based on strong and persistent representation of cue valence in population dynamics.

**Figure 7.**
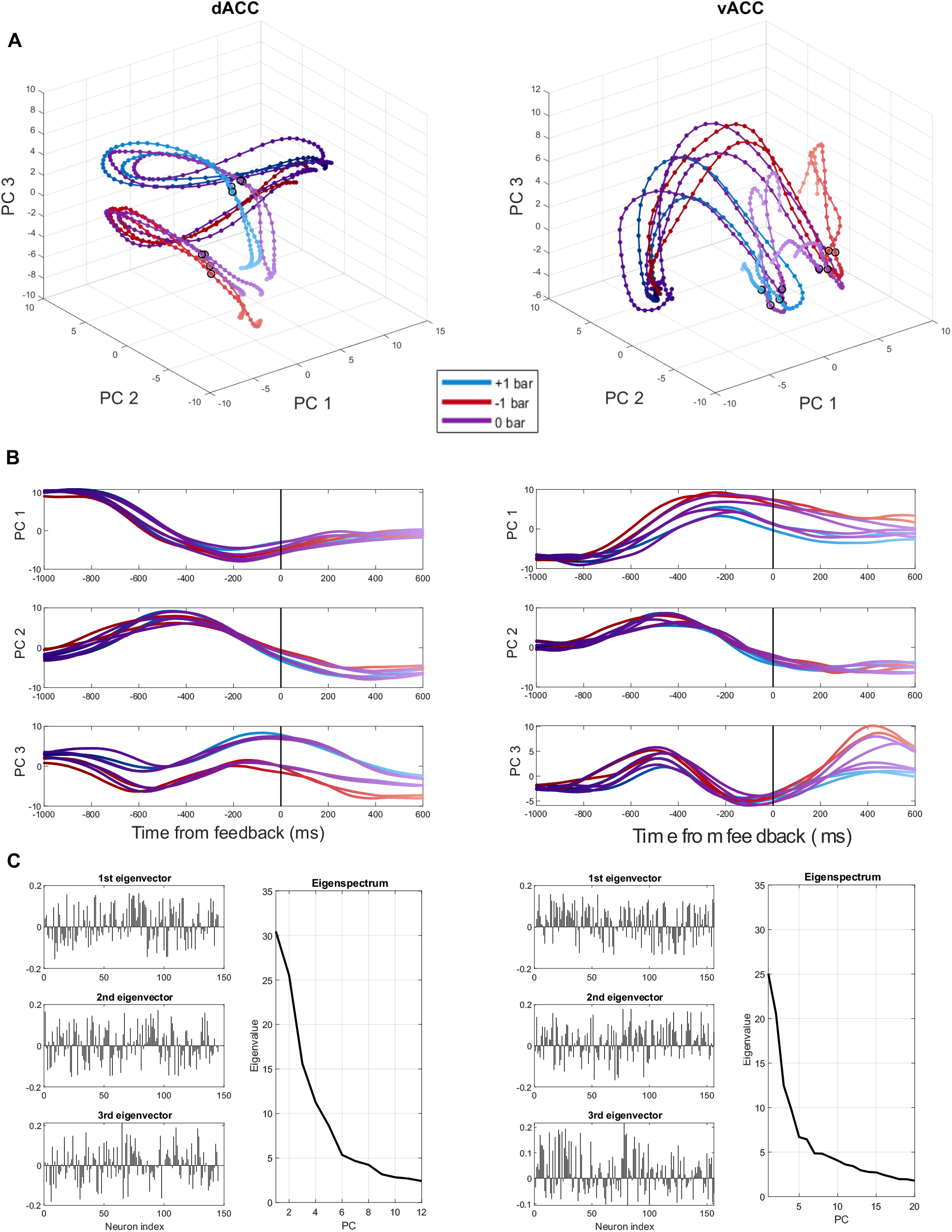
Separation of cue valences is a dominant and stable feature of neural dynamics in ACC. **(A)** PCA trajectories show the first 3 canonical responses through time, across the 8 conditions (one trajectory for each condition), for dACC (left) and vACC (right). Colors correspond to signed magnitude of feedback, and points represent time bins. Darker and lighter points correspond to earlier and later timepoints in the trial, respectively. Feedback onset is marked with black circles. **(B)** Same as **A** but with each PC plotted separately as a function of time. **(C)** Right eigenvectors and eigenvalues of the neuron correlation matrix for dACC (left) and vACC (right). Eigenvector elements define an activation pattern over neurons (a neural mode). The associated canonical response is the activation of this mode at each point in time, across conditions.

## Discussion

The ACC has been implicated in a range of cognitive functions, including adapting behavior to task contexts (17, 41) and linking goals to actions (3, 11, 12, 42, 43). To better understand the role of dACC and vACC in these processes, we investigated how representations of expected outcomes relate to actions and received outcomes. Our results show that single unit responses are heterogeneous, encoding a variety of task-relevant information, including cue valences, response directions and received outcomes. Among valence encoding neurons, there was a tendency toward more encoding of negative information in vACC as in previous studies (34), though our results were inconsistent across subjects. Despite heterogeneity among single neurons, there was clear structure in population activity. In both subregions, representations were primarily organized by a strong and persistent separation of positively and negatively valenced conditions, which indicated the potential for gain or loss on each trial. These results are similar to findings in rodent ACC, where single unit responses are heterogeneous but population activity persistently signals valence throughout the end of a trial and the following inter-trial interval (44). Contrary to other reports (7), however, we did not find interactions between actions and reward contingencies at the level of neuron populations. Representations of cue valence and motor responses had different dynamics and independent influences on population geometries, supporting the notion that ACC treats this information separately. Finally, trial outcomes were primarily coded in a relative manner such that the valence of initial cues provided a context for interpreting the ensuing feedback. Based on this, we suggest that expected outcome encoding in ACC is important for context-setting, and this role explains the strength and persistence of valence encoding in the present task.

### Separation of cue valence and motor response representations

Our results argue against the view that ACC neurons link cognitive processes to goal-directed actions (7, 10, 12, 18, 43, 45). The ACC is strongly connected to the motor system (46, 47), and lesions encompassing dACC and vACC impair the ability to use reward contingencies to guide action selection but not stimulus selection in both monkeys and humans (42, 43, 48). Neurons in ACC encode actions in concert with diverse representations of rewards and other task-relevant information (7, 18, 45, 49 - 55). In particular, some single neurons encode the interaction between actions and outcomes or task contingencies, and this has been interpreted as a mechanism that links these pieces of information (7, 45).

In contrast to this view, we found multiple lines of evidence pointing toward a separation of cue valence and response direction in ACC. In our task, information about reward contingency was delivered as part of compound cues that also instructed a motor response, with potential outcomes and response direction counterbalanced. Both of these variables were encoded by single units, but we did not find evidence that these variables were integrated at the level of population codes. First, cue valence was strongly represented throughout the trial, while response direction was encoded primarily at the time the response was executed, appearing later and disappearing earlier. Second, error trials reduced the decodability of response directions, while leaving cue valence decoding intact. Response coding was also less abstract compared to cue valence. In some cases, cross-condition decoding was even below chance levels, indicating that classifiers trained on one subset of conditions actively misclassified response directions from the held-out conditions. This occurred because the extreme variance along the cue valence dimension caused some decoders tasked with separating response direction to essentially learn cue valences instead. Therefore, generalizability of response information was reduced to support the extreme abstraction of cue valence. Together, these distinctions in population codes depend on parsing the compound cues into separate latent factors, in effect untangling their meaning in order to represent valence and action information separately. Consistent with our results, other neurophysiological studies have emphasized that previously reported action-contingency interactions are relatively uncommon in ACC, and these factors tend to be represented independently (18).

### Valence contexts shape ACC representations

Although our data did not support the idea that ACC encodes interactions of actions and reward contingencies, we did find evidence that ACC is involved in context-dependent behavior. The probabilistic nature of trial outcomes allowed us to dissociate neural responses reflecting the valence of predictive cues from those reflecting the feedback itself. Across multiple analyses, we found that cue valence continued to be represented by neural responses throughout the feedback epoch, even increasing in strength in vACC at that time. We posit that this is because a cue’s valence was important not just for anticipating potential outcomes but also for interpreting the feedback correctly. That is, no change in the size of the reward bar (0 bar condition) could be considered either an optimal or suboptimal outcome, with correct interpretation contingent on the valence of that trial’s cue. From this view, the cue’s valence sets the context for interpreting the outcome.

In line with this, we found that ACC predominantly coded feedback relative to the valence context on each trial. That is, feedback was primarily encoded as either the better or worse possible outcome given the preceding cue, rather than in an absolute format corresponding to a signed magnitude. Notably, encoding the signed magnitude of feedback would be sufficient to optimize task performance by seeking gains and avoiding losses. In addition, encoding cue valence in conjunction with feedback magnitude could allow interpretation of the 0 bar condition. Neither of these strategies require tracking of relative feedback value, yet relative feedback was clearly the dominant format for representing trial outcomes. This is similar to another study that found that single units in ACC coded reward values relative to alternatives available in the trial block, rather than as absolute magnitudes (16). In that case, trial blocks with different reward availability provided contextual information, emphasizing that contexts need not be valenced to impact ACC coding. While valence and context are distinct concepts, the two overlap completely as properties of the cue in our task. In both studies, a relative feedback code can only be computed by taking context into account, and this may explain the prominent encoding of cue valence during the feedback epoch.

### Abstract representations in ACC

Beyond the prevalence and persistence of cue valence signals, we also found that the geometric structure of population responses primarily emphasized an abstract representation of cue valence. Our analyses followed Bernardi *et al*. (25), who proposed that abstraction in neural representations can be identified, in part, by finding how well a decoder trained on one set of stimuli performs when tested on a different stimulus set. If the concept is abstractly represented, the decoding rule should generalize. Such generalization occurs when the variance dedicated to encoding the abstract dimension is large compared to the variance in other dimensions, which is what we found for cue valence. That is, conditions corresponding to different cue valences were widely separated in the neural activity space, while conditions corresponding to the same valence were similar. This resulted in increased reliability of valence representation across various conditions. Moreover, this separation was stable across relevant timescales and accommodated encoding of other task information.

This type of abstraction is consistent with our interpretation that cue valence provided a context signal, insofar as context-dependent behavior entails application of a general principle or rule across various instances. Indeed, abstraction of context information has previously been shown in ACC, as well as in dorsolateral prefrontal cortex and hippocampus, in a contingency reversal task (25). In that study, two other task variables, value and action, were also represented abstractly. In line with this, we found evidence for abstraction of additional variables in this task. The relative feedback signal was represented abstractly in both the dACC and vACC, as was response direction around the time of motor execution. However, the abstract representation of cue valence dominated, likely producing functional clustering in both regions. Variation in population responses due to cue valence was captured by the first PCA principal axis, and removing this variance destroyed most (dACC) or all (vACC) clustering tendencies. Although abstraction can generally be accomplished with random mixing of task variables, the functional clustering we observed is consistent with representational geometries that strongly prioritize the separation of valence contexts, causing this information to appear more prominently across the population (**Figure S7**). Indeed, dACC, which abstracted cue valence more strongly than vACC, also demonstrated stronger functional clustering and higher numbers of neurons selective for cue valence. Altogether, the strong and persistent encoding of valence set this variable apart from the others as a particularly important feature of the task. Integrating abstract representations of additional task variables (i.e., response direction and relative feedback) into the population geometry helps emphasize independent and important concepts among multiple possible combinations of task variables, thus serving as a scaffold for generalizing to new instances.

### Abstraction in ACC suggests linear disentanglement

Here, we operationally defined abstraction as encoding of a concept not dependent on any particular instance thereof (25), as when two different cues with the same valence are represented identically. This type of encoding is also interpretable as a relative invariance to other sources of information. That is, the response directions instructed by two cues of the same valence can differ, but this does not impact valence coding. In machine learning, the ability to represent a task-relevant feature invariant to others in the environment can lead to efficient performance of high-level tasks. For instance, object representations that are invariant to illumination or orientation improve object recognition performance (57). In broader cognitive operations, concept abstraction allows behavior to be more flexible by adapting to different scenarios and transferring to new tasks. Moreover, it enables information to be compressed, which may allow efficient solutions to complex computations (58, 59).

Learning invariant representations can be accomplished by decomposing inputs into separate generative factors, such that each factor’s representation lies in its own subspace that is invariant to transformations defined by the others. This process is referred to as ‘untangling’ or ‘disentangling’ and has been identified as part of the solution to many complex tasks in neuroscience and machine learning (60, 61, 62, 63, 64). For instance, disentangled representations of cue valence and response direction in our task would mean that the transformation of moving in the neural activity space from left to right motor responses should not affect the representation of cue valence, which is consistent with our data. In addition, a simple PCA of trial and time averaged neural activity was remarkably effective for identifying distinct and orthogonal latent dimensions, along which components corresponded to single task factors (cue valence, relative feedback, and to a lesser extent response direction). Crucially, linear combinations of these three leading components also explained the large majority of population variance, with a weaker fourth component capturing a nonlinear combination of cue valence and relative feedback, which is equivalent to the bar change versus no change dichotomy necessary for absolute feedback encoding. Together, these results suggest that condition centroids lie near the vertices of a 3-dimensional hepercuboid in neural activity space, such that transformations for each factor are close to linear and act primarily on separate respective orthogonal subspaces, where each subspace encodes a single latent factor (**Figure S8**).

Since disentangled representations in general need not be defined in linear spaces, the associated transformations need not be linear with respect to the representation’s embedding vector space (here the neural activity space). However, if the transformations are indeed linear, they preserve the structure of the embedding vector space, and the associated representations have been referred to as ‘linearly disentangled’ (60). The theory of abstraction described by Bernardi *et al*. (25) relies on a geometric manifestation of this preservation of structure under linear transformations, namely the preservation of parallelisms, which the CCGP (using linear classifiers) and PS tests assess directly. For instance, a linear decoder can generalize relative feedback across two different cue valences (i.e., [-cue/0bar] vs [-cue/-bar] generalizes to [+cue/+bar] vs [+cue/0bar]) if the negative to positive cue valence transformation entails linear translation along an axis that is orthogonal to a relative feedback axis. In this case, normal vectors to the hyperplanes that separate better and worse outcomes in the two cue valence conditions remain close to parallel (**Figure S8**), suggesting generalization of the linear readout for relative feedback across valence conditions. In our data, we found that such linear separations generalize well across conditions and that vectors linking condition centroids on opposite sides of dichotomies (such as cue valence and relative feedback value) are more parallel than a null model. In addition, the shattering dimensionality for both subjects in both epochs was close to that of a linear model generated based on linear combinations of task factors. Therefore, the possibility of linear disentanglement appears well supported in our data.

## Conclusions

Overall, we found that population activity in dACC and vACC exhibits robust low-dimensional structure in a stimulus-motor mapping task. This complements a growing literature showing that cortical representations of well-learned tasks typically occupy less than the high dimensional potential of the recorded neurons (22 - 25, 65 - 69). These lower dimensional representations are less flexible in that fewer combinations of variables can be read out linearly, which may slow new learning (65), but such representations allow key task variables to be represented in a more robust and generalizable fashion. We present evidence that low dimensional representations in ACC are shaped by a strong and persistent encoding of context. Although there were some differences in the profile of single unit responses in the two subregions, this feature of the population response was consistent. Therefore, we conclude that contextual information is likely a primary driver of population responses in ACC, with additional relevant information integrated into this geometry as needed. These representations likely support context-dependent operations, such as encoding outcomes relative to expected results.

## Materials and Methods

### Subjects and Behavioral Task

Behavioral data have been previously described in (35). Subjects were two male rhesus monkeys (*Macaca mulatta*), M and C, aged 6 and 10 years and weighing approximately 11.0 and 14.5 kg respectively at the time of recording. Subjects sat in a primate chair and viewed a computer screen. Affixed to the front of the chair was a joystick that could be displaced to the left or to the right with minimal force. Cues were presented on the computer screen, behavioral contingencies controlled using Monkey Logic software (70), and eye movements tracked with an infrared camera system (ISCAN). Each subject was chronically implanted with a titanium head positioner and two cylindrical titanium chambers, positioned over target areas in the frontal lobe in each hemisphere. All procedures were performed in accord with the National Institute of Health guidelines and recommendations of the University of California at Berkely Animal Care and Use Committee.

Each recording session consisted of several hundred consecutive trials. At the start of each trial, subjects maintained their gaze within a 1.3° degree radius of a central fixation spot for 650 ms. Immediately following this fixation period, one of four cues appeared on the screen. Cues consisted of images of natural scenes, approximately 2° × 3° in size. Prior to recording, subjects had been trained to move the joystick in a particular direction (left or right) to either earn reward or avoid a loss. Subjects were discouraged from arbitrary responding by penalizing responses within 150 ms of cue presentation with a 5 s time-out but were otherwise free to respond as quickly as they would like. Following a joystick response, subjects received feedback through changes in the length of an onscreen reward bar. The bar was visible at the bottom of the task screen throughout the session and indicated the amount of reward the subject had cached at each point in time. At the end of every block of six completed trials, the subject earned a juice reward proportional in amount to the current length of the bar. The bar was then reset to length = 2 to start the next trial block.

On trials featuring one of the two positively valenced cues (referred to as “positive cues”), a correct response resulted in an increase to the length of the reward bar by 1 increment (+1 bar) and an incorrect response resulted in no change to the bar (0 bar). On trials with negative cues, a correct response resulted in no bar change (0 bar), while an incorrect response resulted in decreased bar length by one increment (−1 bar) (**Figure 1**). The subjects were well trained on the task and performed with high accuracies (35). To ensure sufficient numbers of each of the four possible trial outcomes [+cue/+1 bar], [+cue/+0 bar], [-cue/+0 bar], and [-cue/-1 bar], the 0 bar outcome was delivered on 15% of all positive cue trials, and the -1 bar outcome was delivered on 15% of negative cue trials, regardless of the subject’s response.

### Neural Recording and General Preprocessing

Neural recordings were obtained on a Plexon MAP system. In each session, 4-20 tungsten microelectrodes (FHC, Inc.) were acutely placed in target regions (**Figure S1**). Electrode paths were calculated based on 1.5T MRI scans of each subject’s brain obtained prior to recording. Recorded neurons were not screened for response properties, and therefore represented a random sample in each area. Waveforms were digitized and well-isolated units identified with Offline Sorter (Plexon) and saved for further analysis.

Following spike-sorting and prior to subsequent analyses, we removed any neurons whose mean firing rate across an entire session was less than 1 Hz, as such neurons would not provide sufficient activity to analyze statistically.

### Single Unit Regressions

The selectivity of single neurons was assessed with multiple linear regression models. For each neuron on each trial, we averaged spike counts in 150 ms time bins stepped at 25 ms, with bin centers ranging from 200 ms preceding cue presentation to 650 ms following presentation. For the *i*^*th*^ time bin, we fit an ordinary least squares model:

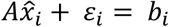

where *b*_*i*_ is the neuron firing rate vector over *m* trials (*m* varies by session), *ε*_*i*_ the error term, 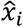 the vector of least squares coefficients, and *A* the *m* × 6 matrix whose columns consist of 5 predictors and an intercept term. These predictors included cue valence and joystick response direction on each trial, coded as 1 for positive valence and leftward response, and -1 for negative valence and rightward response, as well as the interaction of these two terms. In addition, we included two predictors informed by our previous work in which the same behavioral data were analyzed (35), which were log transformations of the size of the reward bar on the current trial and the trial number within a block, both mean-centered at 0. For the purposes of this study, these variables were not analyzed further, but included to improve the accuracy of the regression models.

To assess selectivity of neurons for task variables, we examined the p values for beta coefficients of each predictor. To control Type I errors in the pre-cue epoch (false hits < 0.05 of the population in all bins preceding cue), we imposed a significance level of 0.005 for p values in each bin and deemed significant only those bins that were part of a consecutive series of 3 bins below the 0.005 threshold. Any neuron that passed these criteria during the analysis window (101 to 600 ms following cue and feedback onset) was counted as significant. An overall direction of activation was assigned to significant neurons based on the sign of the mean beta coefficient during this time window.

The coefficient of partial determination (CPD) was computed for the *k*^*th*^ predictor in the *i*^*th*^ time bin as:

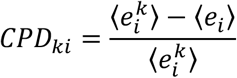

where *e*_*i*_ denotes the vector of residuals for the *i*^*th*^ bin, 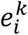 denotes the vector of residuals for the *i*^*th*^ bin with the *k*^*th*^ predictor omitted (reduced model), and ⟨*ν*⟩ = *ν*^*T*^*ν* for a column vector *ν*. The CPD thus assess the strength of a neuron’s encoding of the *k*^*th*^ predictor as the proportion of the firing rate variance left unexplained by the other predictors alone (sum of squared residuals of the reduced model) that is explained by the addition of the *k*^*th*^ predictor. To compare encoding strength between regions, we averaged CPDs within the analysis window for each neuron in the population and compared these mean CPDs between regions using a ranksum test.

During the feedback epoch, we aligned firing rates to the times the reward bar did or would have changed. Similar models assessed coding of relative feedback by replacing cue valence in the predictor matrix. To assess coding of absolute feedback magnitude, we fit an additional model where the predictor matrix included absolute feedback as a signed magnitude rather than cue valence, with all other predictors unchanged. Since the two models had the same number of free parameters, we compared the coefficient of determination (*R*^2^) of the original model (featuring the cue valence predictor) with the *R*^2^ of the model with the absolute feedback predictor for each neuron in each bin. An overall model preference was assigned to each neuron by averaging the difference in *R*^2^ of the two models across bins in the window from 101 ms to 600 ms following feedback presentation, contingent upon that neuron also displaying significant selectivity for the predictor of the better model, as described above. Comparisons based on beta coefficients and CPDs yielded qualitatively similar results.

### Preprocessing for population analyses

Single unit data were aggregated to form pseudopopulations for each subject and each area, and all descriptions of “populations” below refer to pseudopopulations. Here, we excluded any neuron without at least 10 trials in each condition, as smaller trial counts might not support reliable resampling as described below (neuron counts remained the same for both regions in both subjects, except C vACC, where we excluded 6 of 70 neurons; however, this exclusion did not noticeably impact results). Here, we combined neurons from the subjects since we observed good overall agreement between separate population analyses by subject. Spike counts sampled at 1 ms were aligned to the onset of the cue or feedback. Trials from each condition were randomly subsampled to select *n*_*i*_ trials, where *n*_*i*_ is the smallest number of trials from the *i*^*th*^ condition across all sessions. We then generated a *t* × *n* × *p* array, where *t* is the number of timepoints within the analysis window, *n* the sum of the smallest trial counts from each condition, and *p* the number of neurons across all session. We next averaged the data array in 150 ms bins across the first array dimension, stepped at intervals of 25 ms, yielding a *b* × *n* × *p* array, where *b* is the number of 150 ms bins. Activity of each neuron in the array was z-scored across both bins and trials.

### Neural decoding

Linear SVMs were used to classify trials of pseudopopulation responses according to task variables. All task variables were binary, except for absolute feedback, where there were 3 classes. This multiclass classification was implemented using an ensemble method (fitcecoc.m in MATLAB) that combines the performance of 3 binary one-vs-all linear SVMs. SVMs were trained and tested with a resampling strategy as follows. For each time bin, we first randomly split all *n* trials into 10 cross-validation (CV) folds. We then resampled with replacement 100 trials for each of 8 conditions (4 cues x 2 possible outcomes), for each neuron separately (any non-task-related covariance was already destroyed in the initial construction of the pseudopopulation array), yielding an 800 x *p* matrix whose rows are *p*-dimensional single-trial population response vectors. Each of the single-trial vectors was assigned a class label that depended on the variable being decoded (see **Table 2**). For decoding of absolute feedback magnitudes, this resulted in unbalanced class sizes (i.e., there were twice as many 0 bar trials as +1 or -1, since the 0 bar result occurs in 4 of the 8 conditions), so we randomly subsampled the resampled 0 bar trials to match the size of the +1 bar and -1 bar classes. Not performing this subsampling, so that class sizes for the 0-vs-all binary classifier were balanced, yielded qualitatively similar results.

The same resampling strategy was carried out for the held-out data fold, generating another 800 x *p* matrix as a test set. Crucially, single trials were first separated into training and test sets, and resampling was performed separately for these groups so that the same data were never used for both training and testing. Final performance was based on the concatenation of predicted labels from each of the test sets across folds. The entire CV process (over 10 folds) was repeated 100 times for each bin, randomizing the partition of the *n* single trials into folds with each repetition. The final decoding performance for the *i*^*th*^ bin was derived by averaging across the 100 repetitions. Decoding performance timeseries were obtained by performing this CV process in each of the *b n* × *p* slices of the full *b* × *n* × *p* array.

Decoding performance measures included accuracy, balanced accuracy, area under the ROC curve (AUC), and area under the precision-recall curve (AUCPR). Additionally, we assessed decoding performance on each of the classes individually using precision, recall, and the F-measure. We observed good agreement among these calculated metrics. For binary task variables, we report accuracy; for multiclass decoding, we report accuracy as well as the F-measure of the binary one-vs-all decoders for each of the classes. P values were computed by repeating the entire process above, including resampling of training and test sets within each CV iteration, on 1000 data sets in which class labels were randomly permuted (though for each permutation, we performed only one repetition of the entire CV process, rather than 100). P values are derived as: 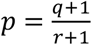, where *q* is the number of random permutations yielding decoding performance equal to or higher than the performance on unpermuted data, and *r* is the number of permutations. The addition of 1 to the numerator and denominator of this ratio is a bias correction for the use of random permutations (71). We set the significance level in each bin as *α* = ⌊0.05/6⌋ = 0.008, where ⌊·⌋ is the floor function, and division by 6 accounts for the fact that a given timepoint can feature in as many as 6 overlapping bins. We note that there is sometimes some small significance prior to median cue onset due to varying latency of motor responses.

### Representational geometry

We assessed measures of representational geometry by generally following previously described methods (25). We first preprocessed the data as above, then resampled trials from each task condition as described in the neural decoding sections, except drawing from all available trials within each condition without first subsampling. For the main results in **Figure 3**, we averaged spike counts in a single time window from 101 to 600 ms following cue or feedback onset. Timeseries results were derived from mean spike counts taken in consecutive, nonoverlapping 100 ms bins. Each neuron’s firing rates were independently z-scored across trials for single window analyses or across trials and time bins for timeseries analyses.

### Cross-Condition Generalization Performance (CCGP)

We first identified sets of conditions that define dichotomies as described in (25). For a set *Z* of *m* integers {1, 2, …, *m*} indexing the task conditions, there are 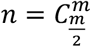 ways to uniquely partition the *m* elements of *Z* into two equal subsets *A* and *B*, each of cardinality 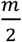. In this calculation, each of the *n* partitions is unique only up to the labels *A* and *B* exchanging places. For example, if one partition has *A* = {2, 4,7, 8} and *B* = {1, 3, 5, 6}, there exists a counterpart partition with *B* = {2, 4,7, 8} and *A* = {1, 3, 5, 6}. The number of completely unique partitions (i.e., not counting the exchange of labels) is thus 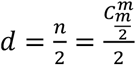. Bernardi *et al*. (25) refer to each such partition as a “dichotomy.” During the cue epoch, there were 4 conditions and *d* = 3 dichotomies. During the feedback epoch, there were 8 conditions and *d* = 35 dichotomies (**Table 2**). In some cases, dichotomies correspond to a binary task variable (e.g., partitioning conditions {1, 3, 5, 7} from conditions {2, 4, 6, 8} corresponds to the relative feedback task variable; see **Table 2**), but most dichotomies result in partitions of conditions not amenable to intuitive interpretation (see **Table S1** for a complete list). The CCGP quantifies how well a separating hyperplane learned by a decoder trained on one subset of conditions within a dichotomy generalizes to the remaining subset. To do this, the test examines all possible ways of choosing subsets of conditions for training and testing for each dichotomy.

More specifically, for a given dichotomy, let *S*_*A,k*_ be the set of all proper subsets of *A* (as defined above) such that each subset *s*_*A,k*_: *s*_*A,k*_ ∈ *S*_*A,k*_ is a set of condition indices of cardinality 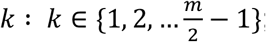; the cardinality of *S*_*A,k*_ is then 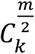. Let *S*_*B,k*_ and *s*_*B,k*_ be defined identically to *S*_*A,k*_ and *s*_*A,k*_, respectively, but based on proper subsets of *B* rather than *A*. For each *s*_*A,k*_, we choose conditions indexed by *s*_*A,k*_ and match them against conditions indexed by each *s*_*B,k*_, i.e., each of the elements of *S*_*B,k*_. For each such pairing, single trials from the matched conditions constitute the training set, with trials from conditions on opposite sides of the dichotomy assigned different class labels. For a given *s*_*A,k*_ and *s*_*B,k*_, the test set consists of trials from the remaining conditions on each side of the dichotomy, i.e., conditions indexed by {*s*_*A,k*_\*A*} and {*s*_*B,k*_\*B*}.

Across all values of *k*, the total number *q* of classifier models trained and tested is given by 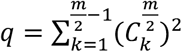, equal to 4 if *m* = 4 during the cue epoch, and 68 if *m* = 8 during the feedback epoch. The CCGP is the average performance across all *q* classifiers. We used linear SVMs as classifiers.

### Parallelism Score (PS)

The PS measures abstraction using a different approach from CCGP that is based on similar geometric ideas (25). Namely, if the optimal separating hyperplane *V*_1_ learned in one set of conditions generalizes well to another set of conditions, whose optimal separating hyperplane is *V*_2_, then the normal vectors *ν*_1_ and *ν*_2_ to *V*_1_ and *V*_2_ respectively, should be close to parallel. To compute the PS, we took the set of condition indices *A* from one side of a dichotomy and put its elements in one-to-one correspondence with each of the elements of the set of condition indices *B* from the other side of the dichotomy. Keeping the order of *A* fixed, we obtained all correspondences with all 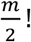 permutations of the elements of *B*. For each such permutation (2 in the cue epoch and 24 in the feedback epoch), we computed the vector difference between the condition centroids—defined by firing rates of the neural population averaged across all single trials of each respective condition—that are matched one-to-one. Thus, for each permutation, there were 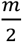 vector differences (2 in the cue epoch and 4 in the feedback epoch).

Next, for each permutation, we considered all 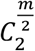 possible ways to pair these vector differences (1 for the cue epoch and 6 for the feedback epoch), and for each pairing calculated the cosine similarity of the pair. We then averaged over all cosine similarities for a given permutation and selected the maximum mean cosine similarity across permutations as the final PS.

### Statistical significance

We used random permutations to generate null distributions for the CCGP and PS tests by independently permuting each column of the predictor matrix. For each permutation, we compute the CCGP and PS for each dichotomy separately. This results in a discrete approximation of one null distribution for each dichotomy, although we observed that these null distributions are in practice often very similar. Two-tailed p values were derived as: *p* = 2 × min(*q*_*L*_, *q*_*R*_), where *q*_*L*_ and *q*_*R*_ are left and right tail probabilities respectively (i.e., the proportion of null models with a CCGP or PS lower than or equal to (respectively, higher than or equal to) that of the neural data; *q*_*L*_ and *q*_*R*_ are corrected for the use of random permutations as described above in *Neural decoding*). Similarly, we used the approximation of the null distribution for each dichotomy to compute 95% confidence intervals around the mean of each null distribution, using the 2.5 and 97.5 percentiles as the lower and upper bounds.

### Shattering Dimensionality (SD)

First, we computed the SD for the recorded neural data. The procedure was the same as that described in the neural decoding section above (with separate resampling within the training and test sets during CV), except that we averaged population activity within a window 101 to 600 ms following cue or feedback onset. We then varied the class labels assigned to trials to decode all possible dichotomies. For example, to decode the cue valence dichotomy, we assigned conditions {1,2,3,4} in **Table 2** a class label of 0 and conditions {5,6,7,8} a class label of 1. To decode the relative outcome dichotomy, we assigned conditions {1,3,5,7} a class label of 0 and conditions {2,4,6,8} a class label of 1. The mean decoding accuracy across all dichotomies is the SD (25).

Next, we adopted a similar approach to Bernardi *et al*. (25) to compare the geometric structure of our data to a linear model (which Bernardi *et al*. term “factorized”). This linear model, in which neural responses are linear combinations of the abstracted task variables, was generated as follows. For *m* task conditions arising as the cartesian product of *k* binary task variables, we modeled condition centroids as vertices of a randomly rotated *k*-dimensional hypercuboid (e.g., as in **Figure 4**) embedded in a *p*-dimensional vector space, where *p* is the number of neurons. During the cue epoch, the relevant task variables are cue valence and response direction (*k* = 2), with relative feedback added during the feedback epoch (*k* = 3). For the *i*^*th*^ condition, we simulated a single-trial point cloud by sampling 100 points from the multivariate Gaussian distribution 𝒩(*μ*_*i*_, 𝕀), where *μ*_*i*_ is the *i*^*th*^ vertex and 𝕀 the identity matrix. The lengths of each of the *k* sides are hyperparameters determining the geometry of the hypercuboid. We initialized the hypercuboid with uniform side lengths of 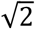 and tuned these *k* hyperparameters to match empirical data by first implementing a rougher-grained pilot grid search, testing *w* = 13 increments to each side length (ranging from -0.5 to 2, stepped by 0.2), exploring a total of *w*^*k*^ distinct configurations for the hypercuboid. For each configuration, we computed the CCGP of the *k* linearly separable dichotomies and used a sum of squared errors loss to select the configuration yielding CCGP values for these task variables most closely empirically observed values. We then conducted a second finer-grained grid search using *ν* increments of 0.02 to each of the side lengths in the neighborhood of the optimal geometry identified in the pilot search, where *ν* is chosen as the number needed to tile the open interval between -0.2 and +0.2 of the optimal increment for each side from the pilot search. No optimal hyperparameters from either search lay at the edge of the grid (which would have triggered a new search with shifted grid values). Finally, we applied our decoding process to the tuned model as for the empirical SD.

### Representational Similarity Analysis

Preprocessing was the same as that described above (*Preprocessing for population analyses*) except that no subsampling by condition was performed here. Briefly, we created a *t* × *c* × *p* array of trial-averaged firing activity where *t* is the number of timepoints at 1 ms resolution, *c* the number of conditions (8, see **Table 2**, Feedback Epoch), and *p* the number of neurons. We then averaged along the first dimension within 150 ms bins stepped at 25 ms to yield a *b* × *c* × *p* firing rate array, where *b* is the number of bins. Each neuron’s activity was z-scored across bins and conditions together. For each bin, we then computed a *c* × *c* condition-wise correlation matrix from the *c* × *p* firing rate matrix. Next, we constructed a series of template correlation matrices, each modeling abstraction of a different task variable. Each template assumes perfect abstraction of a task variable. That is, if the task variable is binary, all conditions that share the same value of this variable (e.g., positive cue valence) will be perfectly correlated with each other and perfectly anticorrelated with conditions having the opposite value of this variable (negative cue valence). We used the following templates (depicted in **Figure 5C**), where *ρ* denotes Pearson correlation:

Template 1: Cue valence, modeling similarity (*ρ* = 11 of conditions featuring the same cue valence, and dissimilarity (*ρ* = −11 of conditions featuring the opposite cue valence.

Template 2: Response direction, modeling similarity (*ρ* = 11 of conditions featuring the same correct response direction and dissimilarity (*ρ* = −11 of conditions featuring the opposite correct response direction.

Template 3: Interaction of cue valence and response direction, modeled as the elementwise product of templates 1 and 2.

Template 4: Relative feedback, modeling similarity (*ρ* = 11 of conditions in which the subject received the feedback with the same relative valence (“better” vs “worse”), and dissimilarity (*ρ* = −11 of conditions in which the subject received feedback oppositely valenced feedback.

Template 5: Absolute feedback, modeling similarity (*ρ* = 11 of conditions in which the subject received the same change (+1, 0, -1) to the reward bar and dissimilarity as follows: all 0 bar conditions are modeled as *ρ* = 0 with respect to all other conditions; -1 and +1 bar conditions are modeled as *ρ* = −1 with respect to each other.

Template 6: 𝕀^m×m^, where *m* equals the number of task conditions, modeling correlations of conditions with themselves, equivalent to unity.

An additional regressor matrix 1^m×m^ was added, whose elements are all unity, allowing an intercept in our regression model.

Template matrices serve as predictors in an ordinary least squares regression that modeled the empirical correlation matrix in each bin as a weighted sum of each template. We assessed the contributions of each template in each bin by computing CPDs as described above (*Single Unit Regressions*). We generated null distributions for CPDs of each template in each bin by randomly permuting the elements of the empirical condition correlation matrix 10,000 times and for each permutation fitting an ordinary least squares model and computing CPDs for each predictor. P values were computed as: 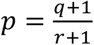, where *q* is the number of random permutations yielding a CPD equal to or higher than the CPD on the unpermuted data, and *r* is the number of random permutations. We imposed in each bin a similar significance threshold of *α* = 0.008 as in the decoding analyses. To control Type I error during the pre-cue epoch, we also stipulated that a bin had to be part of a consecutive series of 4 bins with p < 0.008 to be deemed significant.

### Latent task factors via Principal Components Analysis (PCA)

We performed a Principal Components Analysis (PCA) in order better to understand population structure and task selectivity driven by variance across conditions. First, we formed the *c* × *p* matrix of firing rates averaged across trials and within a window 101 to 600 ms following feedback, where *c* is the number of conditions (8) and *p* the number of neurons. Let 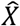 denote this firing rate matrix with columns (neurons) z-scored. We performed PCA by taking the singular value decomposition (SVD): 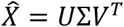. For columns of *U*∑ as principal components (PCs) of 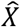, let 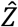 be the matrix whose columns are the top *d* z-scored PCs, with *d* as the minimal number required to explain 90% of the total population variance. For PC variances on the diagonal of 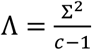, let *H* be the top *d* rows of 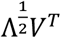. Columns of *H* are loadings, or coordinate vectors with respect to the basis given by columns of 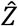, such that we generate a rank *d* approximation of 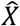 as:

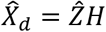

In other words, observed standardized neural responses in columns of 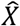 are approximated as linear combinations of *d* standardized principal components, which can be interpreted as latent canonical responses across task conditions. The weights for generating the *i*^*th*^ neuron are elements of the *i*^*th*^ loading vector, stored as the *i*^*th*^ column of *H*. Since *X* was z-scored prior to PCA, these weights are also the respective correlations of the *i*^*th*^ neuron with each of the *d* PCs.

### ePAIRS

To assess functional clustering among ACC neurons, we applied the ePAIRS algorithm, originally proposed by (37) and extended by (38, 40). For each loading vector from the PCA above, the mean angular distance to its *k* nearest neighbors was computed, resulting in a set of *p* mean distances, where *p* is the number of neurons. We used *k* = 3 for our analyses but note that various values of 1, 2, and 9 also give qualitatively similar results. In addition, we used angular distance to emphasize the alignment rather than magnitude in loading space of neurons’ preferred directions. A null model of non-clustered data was constructed following (40), by drawing 1000 random sets of *p* vectors from a multivariate Gaussian distribution with mean and covariance matching the empirical loadings. For each of these 1000 sets of vectors, we then computed mean angular distances for each of the *p* null vectors as for the empirical loadings and used a ranksum test to compare the pooled 1000*p* null distances with the distances from the empirical loadings, yielding a two-sided p value. The effect size *ψ* is computed as:

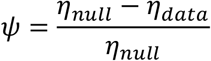

where *η*_*null*_ and *η*_*data*_ are the medians of the pooled null distribution and empirical data respectively. *ψ* > 0 indicates data that display more cluster tendency than expected in the null distribution, while *ψ* < 0 indicates data that are more regularly spaced than expected.

### Dimension removal

To assess the contribution of a given dimension to clustering, we iteratively removed the variance in the original dataset along different principal axes identified with PCA. To do this, we took the SVD of the *c* × *p* firing rates matrix 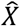 described in the above PCA section: 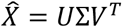, where columns of *V* are principal axes in the neural activity space. We then projected the population activity in 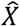 onto a subset, *V*_*r*_, of columns of *V* using the projection matrix *P* = *V*_*r*_*V*_*r*_^*T*^:

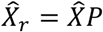

where *V*_*r*_ is equivalent to *V* after dropping columns corresponding to the principal axes along which we would like to remove variance. The projection 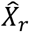 is equivalent to a reconstruction of population activity in the original neural activity space using only the canonical neural responses associated with the columns of 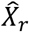. 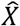 has the same shape as 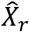, but its rank is decreased by the number of columns of *V* deleted. Note that columns of 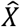 were z-scored again prior to PCA.

### Low-dimensional neural dynamics

To aid in the visualization of neural population dynamics as low-dimensional trajectories, we employed dimensionality reduction using PCA. We began as described in the RSA section above with a *b* × *c* × *p* array of mean firing rates (averaged across trials and within bins), for *b* as the number of 150 ms bins stepped at 25 ms, *c* as the number of conditions (8), and *p* as the number of neurons. Each neuron’s activity was z-scored across bins and conditions together, as in the RSA. Bins and conditions were then vertically concatenated for each neuron to yield a *bc* × *p* firing rate matrix 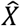. We performed PCA by taking the SVD of 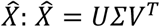. Columns of *UΣ* are principal components (scores) of 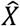 corresponding to latent canonical responses underlying observed neural activity. Columns of *V* are “neural modes” corresponding to canonical patterns of activation over neurons. Each canonical response is the activation across time and conditions of the associated neural mode. By summing the activations of each of the modes, we reconstruct the observed neural population activity (lower case letters denote indexed columns of the matrices denoted by corresponding capital letters):

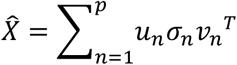

## Acknowledgements

We thank Feng-Kuei Chiang, Pooja Viswanathan, Jessica Overbey, and Peter Rudebeck for discussion and comments on the manuscript. The project was funded by R01DA19028 and P01NS040813 to J.D.W., and R01MH121480 and a grant from the Hilda and Preston Davis Foundation to E.L.R.

## Data Availability

All data, metadata, and custom analysis code related to this study will be uploaded to a DOI-issuing data repository prior to publication. Custom analysis code is currently available at: https://github.com/jonathan-chien/ACC-abstract-representational-geometry.

## Notes

### Competing Interest Statement

The authors have declared no competing interest.

### Summary of Updates

Edits to the text and Figure 6 for clarity. New supplemental figures elaborate on results.

