## Supplemental Information for "Abstraction of reward context facilitates relative reward coding in neural populations of the anterior cingulate cortex"

### Abstraction of reward context facilitates relative reward coding in dorsal and ventral anterior cingulate cortex

**Supplementary Figure S1.**

Target regions and electrode tracks.

**Supplementary Figure S2.**

Single neuron encoding of instructed response directions.

**Supplementary Figure S3**.

Time courses of valence encoding

**Supplementary Figure S4.**

Proportions of single units displaying linear and nonlinear mixed selectivity.

**Supplementary Figure S5.**

Abstract geometry schematic related to Figure 4C.

**Supplementary Figure S6.**

CCGP of the tuned linear model vs empirical data.

**Supplementary Figure S7.**

Simulated and empirical data showing influence of uneven abstraction on loading weights.

**Supplementary Figure S8.**

Abstract geometry schematic related to linear disentanglement.

**Supplementary Table 1.**

Lists of all dichotomies during the cue and feedback epochs.

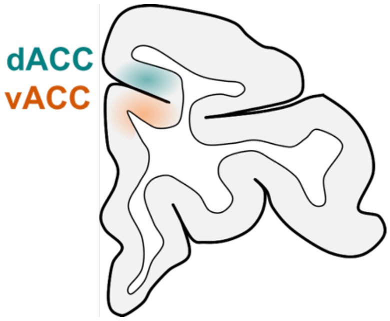


**Figure S1.** Target regions for dACC and vACC. The schematic shows a coronal section of one hemisphere, approximately at the middle of the recording chambers. dACC is shaded in teal, vACC is shaded in orange. Note that vACC targets included the ventral bank of the cingulate sulcus, as well as the most dorsal portion of the cingulate gyrus. Subject M was recorded in the left hemisphere, subject C was recorded in the right.

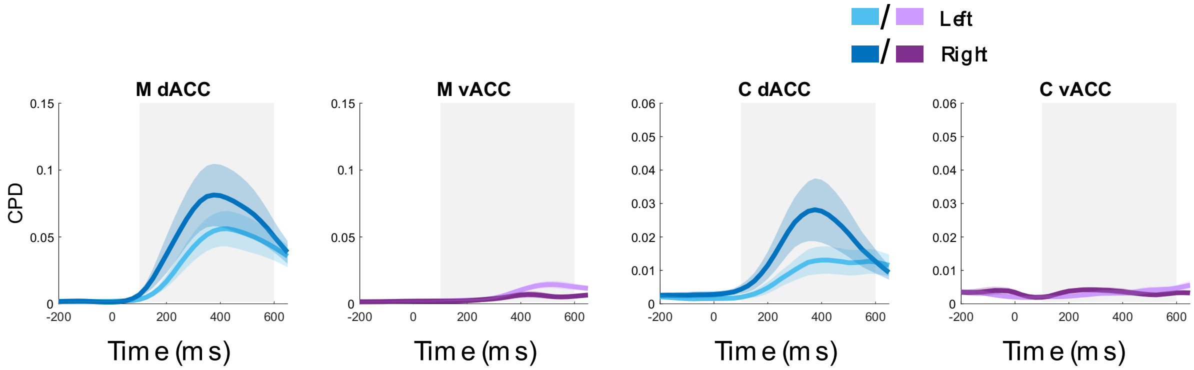


**Figure S2:** Single neuron encoding of instructed response directions. Mean coefficients of partial determination (CPDs) across neurons for the response direction predictor as a function of time, from the same regression model in main text **Figure 2 A & B**. Blues = dACC, Purples = vACC, Light colors = neurons preferring left, Dark colors = neurons preferring right. Time 0 = cue onset, gray = cue epoch. To compare encoding strength between regions, we averaged CPDs within the analysis window for each neuron in the population (differently from the procedure to generate the timeseries plots) and compared these mean CPDs using a Wilcoxon ranksum test. Response encoding was more prominent in dACC in both subjects. In Subject M, there was a significant difference between directions in vACC ($W = 2978$, $p = 0.01$) but not in dACC ($W = 1992$, $p = 0.34$). In Subject C, neither region displayed a significant difference ($W = 927$, $p = 0.52$ and $W = 1227$, $p = 0.15$ for dACC and vACC, respectively).

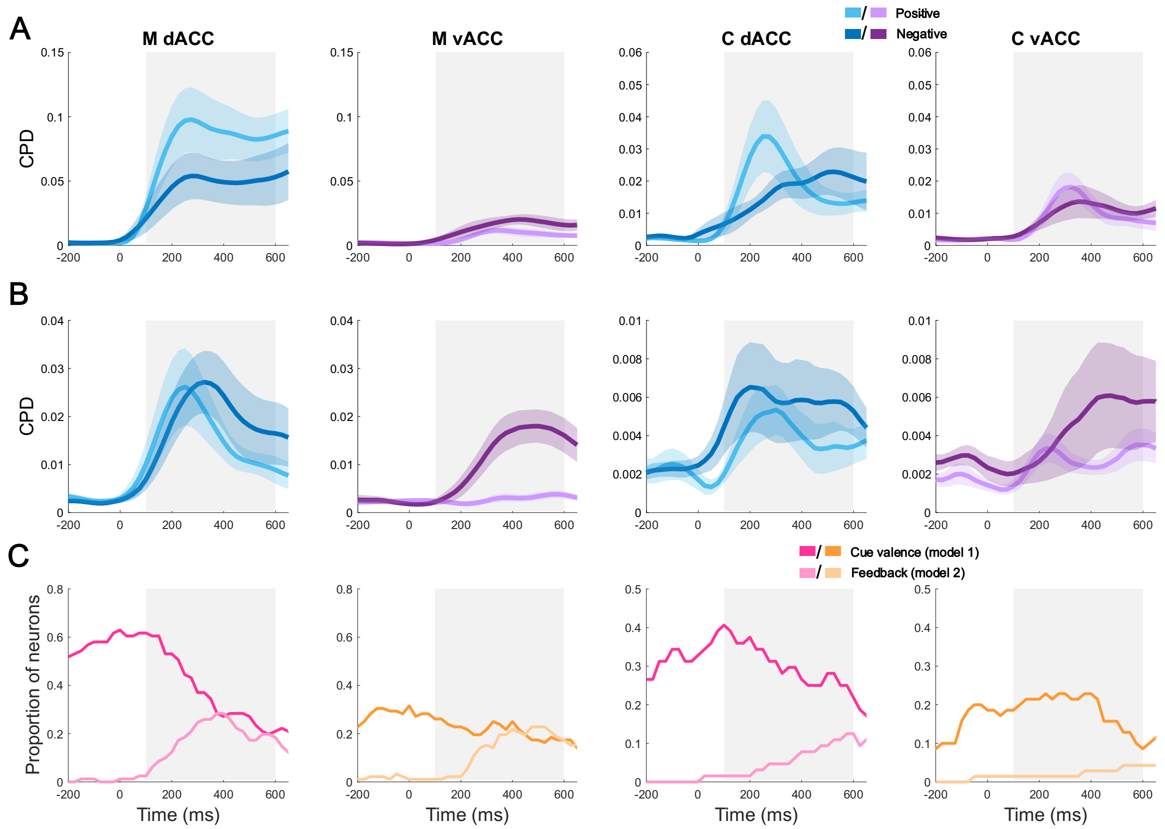


**Figure S3.** Time courses of valence encoding. To further assess valence encoding in the two regions, we considered the contribution of valence to variance in single unit activity, regardless of count-based significance thresholds. In other words, even if the two areas did not differ in the proportion of neurons defined as selective, their responses could still differ in strength. To assess this, we computed the CPD, which is the marginal contribution of one factor toward explaining trial-by-trial variability in firing (**Methods**). **(A)** Mean (± SEM) CPD for cue valence across all single units in dACC (blues) and vACC (purples), for both subjects. Darker colors = larger response to negative cues, lighter colors = positive cues. Time 0 = cue onset, gray = cue epoch. To compare encoding strength between regions, we averaged CPDs within the analysis window for each neuron in the population (differently from the procedure to generate the timeseries plots) and compared these mean CPDs between regions using a ranksum test. There were slightly larger average CPDs for positive compared to negative-responsive neurons (categorized by the beta coefficient sign) in dACC in subject M (Wilcoxon rank-sum, $p = 0.039$, $W = 1859$, effect size = 0.014), but no differences in subject C or in vACC in either subject (Wilcoxon rank-sum tests, all $p > 0.3$). **(B)** Same as **(A)**, but for the relative feedback predictor during the feedback epoch. Time 0 = feedback onset, gray = feedback epoch. Encoding strength was stronger for worse outcomes (negative relative feedback) in vACC in subject M (Wilcoxon rank-sum, $p = 3.23 \times 10^{-4}$, $W = 1313$, effect size = -0.0033), but not C (Wilcoxon rank-sum, $p = 0.79$, $W = 1159$, effect size = $-7.58 \times 10^{-5}$), and there were no differences in either subject in dACC (Wilcoxon rank-sum, both $p>0.3$). **(C)** During each bin, neuron activity was marked as better predicted by a model with a cue valence predictor (model 1, darker colors) or one with an absolute feedback predictor (model 2, lighter colors). Plots show the proportion of neurons (out of the entire population) assigned to each model that were also selective for the respective regressor. Pinks = dACC, oranges = vACC. After the onset of feedback, cue valence encoding decreased somewhat but continued to be encoded throughout the remainder of the trial. At the same time, absolute feedback encoding increased in prevalence, particularly in subject M and less so in subject C.

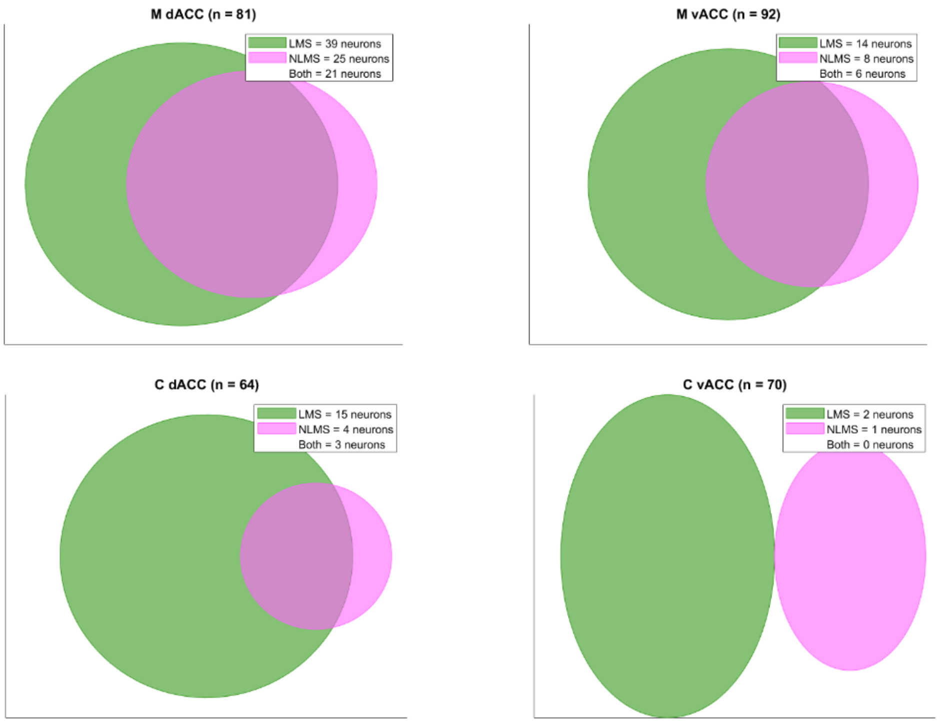


**Figure S4**. Proportions of single units displaying linear and nonlinear mixed selectivity. Panel shows proportions of neurons in each subject and each area selective for both cue valence and response direction during the cue epoch. We considered a unit as displaying linear mixed selectivity (LMS) if our ordinary least squares regression model found that both valence and direction explained significant variance in its firing rate. We considered units as displaying nonlinear mixed selectivity (NLMS) if the interaction term between valence and direction was significant. In subject M, 48% of all dACC units displayed LMS, 31% displayed NLMS, and 26% displayed both. In M, vACC, 15% of all units displayed LMS, 9% displayed NLMS, and 7% displayed both. In subject C in dACC, 23% of all units displayed LMS, 6% displayed NLMS and 5% displayed both. In C vACC, 3% of all units displayed LMS and 1% displayed NLMS, and none displayed both. We also examined mixed selectivity to relative feedback valence and response direction during the feedback epoch (not shown), but found very few neurons encoded both types of information (Subject M, dACC: 8 units with LMS, 1 with NLMS, 0 with both; vACC: 5 units with LMS, 6 with NLMS, 2 with both. Subject C, dACC: 3 units with LMS, 2 with NLMS, 1 with both; vACC: 1 neuron with LMS, 2 with NLMS, 0 with both).

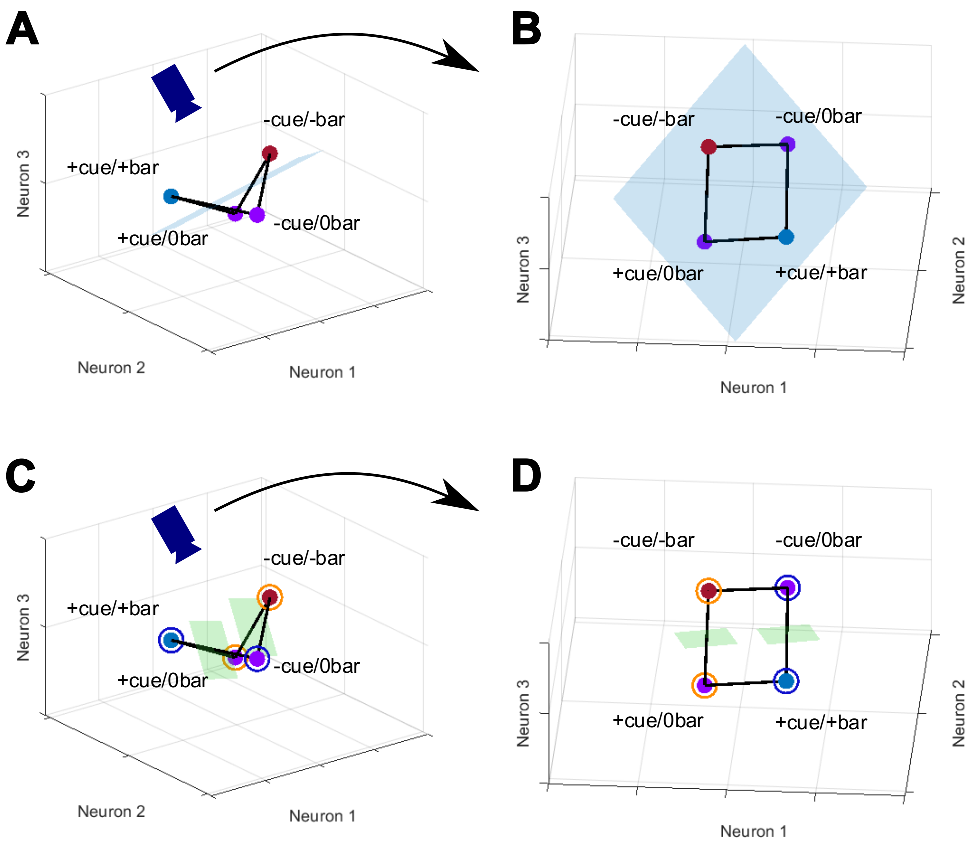


**Figure S5. (A)** Reproduction of **Figure 4C**. **(B)** Same as **A** but viewed from the angle depicted by the camera. **(C)** Training and testing hyperplanes (in green) for assessing CCGP of the dichotomy corresponding to no-bar-change (i.e., 0 bar, purple centroids) vs. bar-change (i.e., +/-bar, blue and red centroids) conditions. **(D)** Same as **C**, but from the angle depicted by the camera. Note that training a decoder to classify no-change vs. change conditions based on the [-cue/0bar] vs. [+cue/+bar] training set (circled in blue) would result in learning the hyperplane (in green) on the right. Using this same hyperplane to classify no-change vs. change conditions in the [+cue/0bar] vs. [-cue/-bar] test set (circled in orange) on the left results in active misclassification (i.e., below chance performance), since the position of purple centroids relative to the hyperplane is reversed between training and testing. The same holds for the other 3 training and testing possibilities: training on [+cue/0bar] vs [-cue/-bar] and testing on [-cue/0bar] vs [+cue/+bar]; training on [+cue/0bar] vs [+cue/+bar] and testing on [-cue/0bar] vs. [-cue/-bar]; training on [-cue/-bar] vs [-cue/0bar] and testing on [+cue/0bar] vs [+cue/+bar]. Thus, though linear separation of this no-feedback vs feedback dichotomy is still possible as shown by the blue hyperplane in **A**, its CCGP will be theoretically 0, as its generalizability is sacrificed (note the minimal variance along this no-change vs change dimension) to accommodate abstract, generalizable representation of the other two dichotomies here (cue valence and relative feedback, respectively).

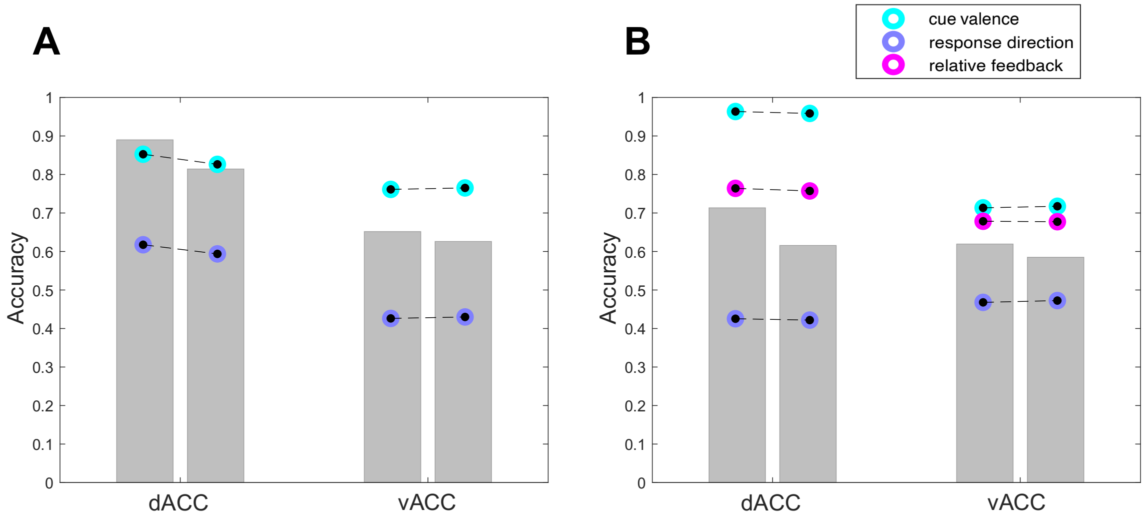


**Figure S6**. **(A)** For each region, the left bar is the shattering dimensionality (SD) of the empirical data over the 3 cue epoch dichotomies (see **Table S1**), while the right bar is the SD of the tuned linear model. Points circled in color are the CCGPs for cue valence and response direction in the empirical data (left) vs the tuned linear model (right). Sum of squared errors between empirical and tuned CCGPs for the 2 highlighted dichotomies is 6.63 x $10^{-4}$ in dACC and 1.41 x $10^{-5}$ in vACC. **(B)** Same as **A** but for the feedback epoch, with 35 dichotomies (see **Table S1**) and tuning performed based on the cue valence, response direction, and relative feedback dichotomies. Sum of squared errors between empirical and tuned CCGPs for the 3 highlighted dichotomies is 4.24 x $10^{-5}$ in dACC and 1.74 x $10^{-5}$ in vACC. Note that even though CCGPs of the empirical vs. linear model are matched for a given dichotomy, the decoding accuracy for that dichotomy in the empirical data can still be higher than in the linear model (e.g., compare decoding accuracy of relative feedback and response direction for the empirical data vs linear model in **Figure 3J**). This is likely because nonlinearity in the empirical data compromises parallelism, which generally decreases CCGP, but this effect can be compensated for by increasing the variance between opposite sides of the dichotomy, resulting in higher decoding accuracy compared to the linear model. Note as well that decodability and CCGP are generally not the same, with sufficient conditions for the former being less stringent than those for the latter.

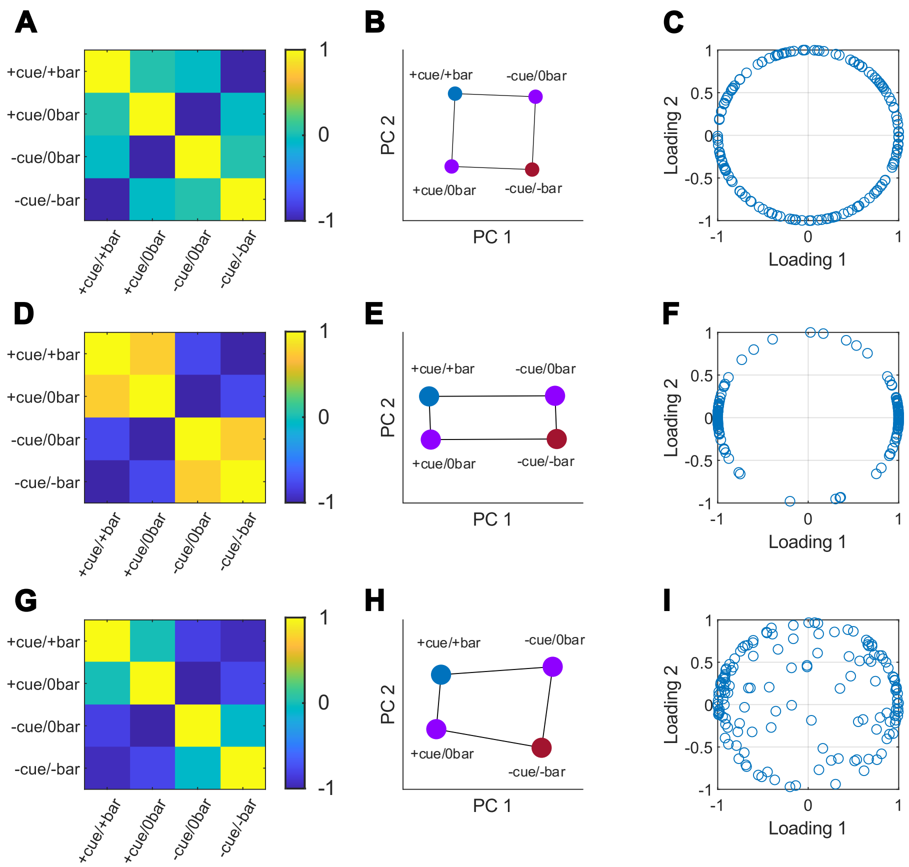


**Figure S7**. Condition-wise population responses for real and simulated data. **(A-C)** Simulated neural data consisting of 145 neurons in 4 conditions corresponding to possible valence-outcome combinations. **(B)** This simulation shows a fully factorized population structure, where neural responses are random linear combinations of cue valence and relative feedback value, recoverable by PCA. **(C)** This results in randomly distributed PC loadings. A neuron $x$ has a 2-dimensional loading vector $h$ whose elements are respective correlations of $x$ with the two PCs (**Methods**). Since $x$ and any given PC both have 0 mean, their correlation is equivalent to their cosine similarity. Let $\theta$ be the angle between $x$ and PC1. PCs are chosen orthogonal, so the angle between $x$ and PC 2 is $\frac{\pi}{2} - \theta$. Since cosine is even, $\cos\left(\frac{\pi}{2} - \theta\right) = \cos\left(-\left(\theta - \frac{\pi}{2}\right)\right) = \cos\left(\theta - \frac{\pi}{2}\right) = \sin\theta$. Thus, the space of all possible loadings when normalized data are linear combinations of two latent variables is parameterized by the unit circle: $h = \begin{bmatrix}\cos\theta \\ \sin\theta\end{bmatrix}$. In general, $n$ latent variables result in loadings on a unit ($n$-1)-sphere. (**D-F**) Same as **A**, but if cue valence (PC 1) variance >> relative feedback (PC 2), which has nonzero variance. In this case, the population displays relative insensitivity to relative feedback and abstraction of and selectivity for cue valence. Loadings are then clustered near a 0-sphere (if relative feedback variance = 0, they lie exactly on the 0-sphere), indicating that all neurons are strongly correlated or anticorrelated with a canonical cue valence response and mostly invariant to relative feedback, even though the geometry is still randomly rotated in neural activity space. **(G-I)**. Same as **A-C, D-F** but for empirical spiking data recorded in dACC, averaged across trials and time from 101 to 600 ms post feedback. Note that these data emphasize the separation of cue valence, but still contain enough nonlinearities (a canonical response encoding bar change vs no-change as a nonlinear combination of cue valence and relative feedback, captured by a third PC (see **Figure 4B and 6A-C**)) to permit absolute feedback decoding. Cluster structure near the first loading axis **(I)** indicates that more neurons are correlated with PC1 (cue valence) than expected if variables are abstracted with equal strength. This corresponds to the dominant abstraction of cue valence in this population. If there are more than 2 latent variables, this binary cluster structure can be discerned as long as one dimension dominates strongly over the rest. If this dominant dimension (e.g., cue valence) is removed, one of the remaining dimensions (e.g., relative feedback) may then drive cluster structure, provided it is relatively dominant over the other remaining dimensions.

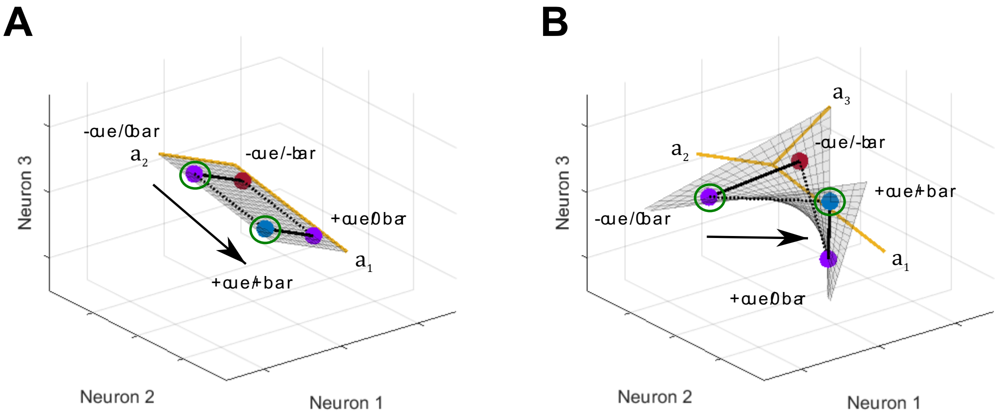


**Figure S8. (A)** Schematic of a linearly disentangled neural representation resulting from linear combinations of cue valence and relative feedback. The representation lies in a 2D vector space for which $a_1$ and $a_2$ (PCA principal axes capturing cue valence and relative feedback, respectively) form a basis. A linear decoder can robustly generalize relative feedback value across two different cue valences since the linear transformation (black arrow) from negative to positive cue valence (moving the circled purple point to the circled blue point) acts along only $a_1$ ($a_2$ is left as is). Note that normal vectors (bold black) to relative feedback separating hyperplanes in the two cue valence conditions are parallel. **(B)** Same as **A**, but neural responses incorporate a nonlinear combination of cue valence and relative feedback. Note that the normal vectors (bold black) from **A** are no longer parallel. The negative to positive cue valence transformation now occurs on a nonlinear manifold embedded in a 3D vector space for which principal axes $a_1$, $a_2$, and $a_3$ (capturing cue valence, relative feedback, and bar change vs no change, respectively) form a basis. The corresponding linear transformation (black arrow) from the circled purple point (negative valence) to the circled blue point (positive valence) acts along both $a_1$ and $a_3$ (the linking vector difference (dotted line) has nonzero components with respect to both $a_1$ and $a_3$). I.e., changing cue valence requires a nontrivial change along the bar change axis as well, so these two factors are linearly entangled, leading to worse generalization by a linear decoder in the presence of noise. If the vector difference linking two condition centroids has nonzero components along both cue valence and bar change axes, a cue valence decoder and a bar change decoder will both learn the same dichotomy, namely whichever dichotomy contributes the larger component to this vector difference. This underlies the below-chance CCGP for all training and testing subsets of the bar change vs no change dichotomy in **Figure S5**, as well as the below-chance CCGP for training and testing subsets across the diagonal for the response direction dichotomy in **Figure 4E**. In general, for representations to be linearly disentangled, each transformation must be linear and act nontrivially only on the subspace corresponding to a single task factor, as in **A**. The representation in **B** may still be considered disentangled with respect to nonlinear transformations that do not preserve the structure of the vector space, though this requires more complicated nonlinear readouts. Real neural populations may favor simpler linear readouts but still incorporate weak and specific nonlinearities (e.g., if variance along $a_3$ is **B** is small), thus exploiting additional linear separations while remaining close to being linearly disentangled.

### Supplementary Table 1

List of all dichotomies, with rows corresponding to dichotomies and columns to task conditions (See **Table 2** for explanation of condition abbreviations). For $m$ total conditions, the first $\frac{m}{2}$ conditions in each row constitute one side of the dichotomy and the remaining conditions the other. Dichotomies corresponding to interpretable task variables are set in boldface and color coded.

#### Cue epoch dichotomies (4 conditions, 3 dichotomies)

| | 1 | 2 | 3 | 4 |
|---|---|---|---|---|
| **Cue valence** | **+cue/L** | **+cue/R** | **-cue/L** | **-cue/R** |
| **Response direction** | **+cue/L** | **-cue/L** | **+cue/R** | **-cue/R** |
| | +cue/L | -cue/R | +cue/R | -cue/L |

#### Feedback epoch dichotomies (8 conditions, 35 dichotomies)

| 1 | 2 | 3 | 4 | 5 | 6 | 7 | 8 |
|---|---|---|---|---|---|---|---|
| **+cue/L/+bar** | **+cue/L/0bar** | **+cue/R/+bar** | **+cue/R/0bar** | **-cue/L/0bar** | **-cue/L/-bar** | **-cue/R/0bar** | **-cue/R/-bar** |
| +cue/L/+bar | +cue/L/0bar | +cue/R/+bar | -cue/L/0bar | +cue/R/0bar | -cue/L/-bar | -cue/R/0bar | -cue/R/-bar |
| +cue/L/+bar | +cue/L/0bar | +cue/R/+bar | -cue/L/-bar | +cue/R/0bar | -cue/L/0bar | -cue/R/0bar | -cue/R/-bar |
| +cue/L/+bar | +cue/L/0bar | +cue/R/+bar | -cue/R/0bar | +cue/R/0bar | -cue/L/0bar | -cue/L/-bar | -cue/R/-bar |
| +cue/L/+bar | +cue/L/0bar | +cue/R/+bar | -cue/R/-bar | +cue/R/0bar | -cue/L/0bar | -cue/L/-bar | -cue/R/0bar |
| +cue/L/+bar | +cue/L/0bar | +cue/R/0bar | -cue/L/0bar | +cue/R/+bar | -cue/L/-bar | -cue/R/0bar | -cue/R/-bar |
| +cue/L/+bar | +cue/L/0bar | +cue/R/0bar | -cue/L/-bar | +cue/R/+bar | -cue/L/0bar | -cue/R/0bar | -cue/R/-bar |
| +cue/L/+bar | +cue/L/0bar | +cue/R/0bar | -cue/R/0bar | +cue/R/+bar | -cue/L/0bar | -cue/L/-bar | -cue/R/-bar |
| +cue/L/+bar | +cue/L/0bar | +cue/R/0bar | -cue/R/-bar | +cue/R/+bar | -cue/L/0bar | -cue/L/-bar | -cue/R/0bar |
| **+cue/L/+bar** | **+cue/L/0bar** | **-cue/L/0bar** | **-cue/L/-bar** | **+cue/R/+bar** | **+cue/R/0bar** | **-cue/R/0bar** | **-cue/R/-bar** |
| +cue/L/+bar | +cue/L/0bar | -cue/L/0bar | -cue/R/0bar | +cue/R/+bar | +cue/R/0bar | -cue/L/-bar | -cue/R/-bar |
| +cue/L/+bar | +cue/L/0bar | -cue/L/0bar | -cue/R/-bar | +cue/R/+bar | +cue/R/0bar | -cue/L/-bar | -cue/R/0bar |
| +cue/L/+bar | +cue/L/0bar | -cue/L/-bar | -cue/R/0bar | +cue/R/+bar | +cue/R/0bar | -cue/L/0bar | -cue/R/-bar |
| +cue/L/+bar | +cue/L/0bar | -cue/L/-bar | -cue/R/-bar | +cue/R/+bar | +cue/R/0bar | -cue/L/0bar | -cue/R/0bar |
| +cue/L/+bar | +cue/L/0bar | -cue/R/0bar | -cue/R/-bar | +cue/R/+bar | +cue/R/0bar | -cue/L/0bar | -cue/L/-bar |
| +cue/L/+bar | +cue/R/+bar | +cue/R/0bar | -cue/L/0bar | +cue/L/0bar | -cue/L/-bar | -cue/R/0bar | -cue/R/-bar |
| +cue/L/+bar | +cue/R/+bar | +cue/R/0bar | -cue/L/-bar | +cue/L/0bar | -cue/L/0bar | -cue/R/0bar | -cue/R/-bar |
| +cue/L/+bar | +cue/R/+bar | +cue/R/0bar | -cue/R/0bar | +cue/L/0bar | -cue/L/0bar | -cue/L/-bar | -cue/R/-bar |
| +cue/L/+bar | +cue/R/+bar | +cue/R/0bar | -cue/R/-bar | +cue/L/0bar | -cue/L/0bar | -cue/L/-bar | -cue/R/0bar |
| +cue/L/+bar | +cue/R/+bar | -cue/L/0bar | -cue/L/-bar | +cue/L/0bar | +cue/R/0bar | -cue/R/0bar | -cue/R/-bar |
| **+cue/L/+bar** | **+cue/R/+bar** | **-cue/L/0bar** | **-cue/R/0bar** | **+cue/L/0bar** | **+cue/R/0bar** | **-cue/L/-bar** | **-cue/R/-bar** |
| +cue/L/+bar | +cue/R/+bar | -cue/L/0bar | -cue/R/-bar | +cue/L/0bar | +cue/R/0bar | -cue/L/-bar | -cue/R/0bar |
| +cue/L/+bar | +cue/R/+bar | -cue/L/-bar | -cue/R/0bar | +cue/L/0bar | +cue/R/0bar | -cue/L/0bar | -cue/R/-bar |
| **+cue/L/+bar** | **+cue/R/+bar** | **-cue/L/-bar** | **-cue/R/-bar** | **+cue/L/0bar** | **+cue/R/0bar** | **-cue/L/0bar** | **-cue/R/0bar** |
| +cue/L/+bar | +cue/R/+bar | -cue/R/0bar | -cue/R/-bar | +cue/L/0bar | +cue/R/0bar | -cue/L/0bar | -cue/L/-bar |
| +cue/L/+bar | +cue/R/0bar | -cue/L/0bar | -cue/L/-bar | +cue/L/0bar | +cue/R/+bar | -cue/R/0bar | -cue/R/-bar |
| +cue/L/+bar | +cue/R/0bar | -cue/L/0bar | -cue/R/0bar | +cue/L/0bar | +cue/R/+bar | -cue/L/-bar | -cue/R/-bar |
| +cue/L/+bar | +cue/R/0bar | -cue/L/0bar | -cue/R/-bar | +cue/L/0bar | +cue/R/+bar | -cue/L/-bar | -cue/R/0bar |
| +cue/L/+bar | +cue/R/0bar | -cue/L/-bar | -cue/R/0bar | +cue/L/0bar | +cue/R/+bar | -cue/L/0bar | -cue/R/-bar |
| +cue/L/+bar | +cue/R/0bar | -cue/L/-bar | -cue/R/-bar | +cue/L/0bar | +cue/R/+bar | -cue/L/0bar | -cue/R/0bar |
| +cue/L/+bar | +cue/R/0bar | -cue/R/0bar | -cue/R/-bar | +cue/L/0bar | +cue/R/+bar | -cue/L/0bar | -cue/L/-bar |
| +cue/L/+bar | -cue/L/0bar | -cue/L/-bar | -cue/R/0bar | +cue/L/0bar | +cue/R/+bar | +cue/R/0bar | -cue/R/-bar |
| +cue/L/+bar | -cue/L/0bar | -cue/L/-bar | -cue/R/-bar | +cue/L/0bar | +cue/R/+bar | +cue/R/0bar | -cue/R/0bar |
| +cue/L/+bar | -cue/L/0bar | -cue/R/0bar | -cue/R/-bar | +cue/L/0bar | +cue/R/+bar | +cue/R/0bar | -cue/L/-bar |
| +cue/L/+bar | -cue/L/-bar | -cue/R/0bar | -cue/R/-bar | +cue/L/0bar | +cue/R/+bar | +cue/R/0bar | -cue/L/0bar |

**Cue valence**
**Response direction**
**Relative feedback**
**Bar change vs no change**
